# T Cells Tear Apart Confining Extracellular Matrix Via a Breaststroke-like Motion to Generate Migration Paths

**DOI:** 10.1101/2025.10.03.680352

**Authors:** Byunghang Ha, Peter Xie, Benjamin Johns, Cole Allan, Maria Korah, Daniel Delitto, Paul L. Bollyky, Natalie Torok, Ovijit Chaudhuri

**Affiliations:** Department of Mechanical Engineering, Stanford University, Stanford CA, 94305, USA; Department of Internal Medicine Gastroenterology and Hepatology, Stanford University, Stanford CA, 94305, and VA Palo Alto, USA; Department of Surgery, Stanford University, Stanford CA, 94305, USA; Department of Internal Medicine, Infectious Diseases, Stanford University, Stanford CA, 94305, USA; Chemistry, Engineering, and Medicine for Human Health, Stanford University, Stanford, CA, USA

## Abstract

T cells adeptly migrate through soft tissues to target aberrant cells and regulate immunity. However, how they establish migration paths in confining nanoporous extracellular matrices (ECMs), and why they often fail to do so in dense ECMs that occur during fibrosis and around tumors, remain unclear. Here, we studied T cell migration in confining collagen-rich hydrogels spanning a range of stiffness, viscoelasticity, mechanical plasticity, and shear strength. Strikingly, only shear strength—the stress required for material failure—correlated strongly with migration, challenging the long-held focus on stiffness and pore size in cell motility. During migration, T cells extend actin-rich, finger-like protrusions into the ECM, which then undergo divergent breaststroke-like motion. Thus, T cells tear apart confining matrices using breaststroke-like motion to generate migration paths.

## Main Text

T cells continuously migrate through soft tissue extracellular matrix (ECM) to identify and eliminate cancerous cells (*1, 2*) (**Fig 1A**). In healthy soft tissues, the ECM is typically comprised of a type-1 collagen (col-1) rich fibrillar matrix, which plays a key role in governing overall tissue structure and mechanics. Numerous other matrix components including elastin, hyaluronic acid (HA), proteoglycans, and various other components also comprise the healthy ECM (*3*). While the collagen network pore size can be on the micron-scale in healthy tissues, these other components fill the space between the collagen fibers and reduce the overall matrix pore size likely to the nanometers scale, though exact measurements of overall matrix pore size are technically challenging (*4, 5*). However, during cancer progression, the matrix becomes much denser and more disorganized, often with much higher levels of col-1 and HA, leading to profound changes in the mechanical properties of cancerous tissues (*6, 7*). While previous studies have focused on the increase in stiffness that often accompanies tumor progression, recent studies point to changes in other mechanical properties, such as viscoelasticity, referring to viscous characteristics and the time-dependence of the mechanical response, and matrix mechanical plasticity, or the ability of the matrix to sustain permanent deformations (*8, 9*). Changes in ECM properties are accompanied by reduced T cell infiltration in “cold” tumors and diminished efficacy of immunotherapies that target T cells (*10*). The extent to which matrix mechanical properties in nanoporous, confining environments govern T cell migration remains unclear, but likely represents a key barrier underlying immunotherapy resistance.

**Fig. 1.**
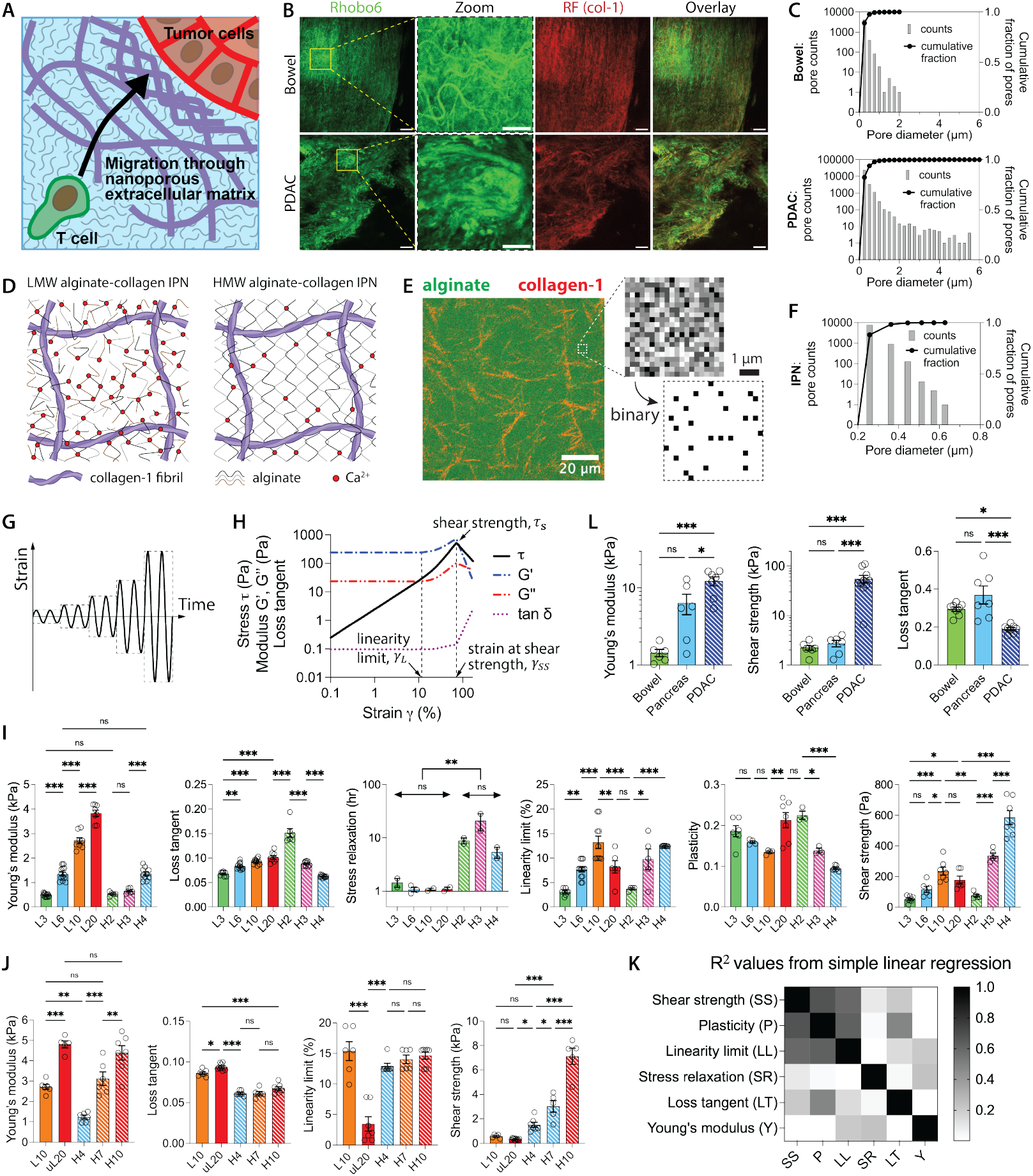
Collagen-rich, nanoporous hydrogel platforms with a range of stiffness, viscoelasticity, mechanical plasticity, and shear strength relevant to both normal and tumor stromal tissues. (A) Schematic illustrating T cell migration through confining ECM to target tumor cells. (B) Representative confocal micrographs of normal bowel (N=3) and PDAC (N=9) tissues reveals a dense ECM architecture composed of glycan-bearing polymers (green) and collagen-1 (red). Scale bars: 50 µm (main) and 20 µm (inset). (C) Pore size analysis of images in (B). The x-axis is limited to 6 μm to better visualize submicron pore sizes. Refer to Fig. S1 for all 12 tissue images and analyses with full x-axis range. (D) Schematic of the IPNs made with low molecular weight (LMW, left) and high molecular weight (HMW, right) alginate. (E) Micrograph showing an IPN architecture, including collagen imaged via reflectance microscopy overlaid with fluorescent alginate. Top inset showing zoomed-in view of alginate fluorescence and bottom inset, the binary processed of the top, showing pores as below-threshold fluorescence pixels. (F) Pore size analysis of the image in (E). (G) Schematic illustrating stepwise increments of applied strain during amplitude sweeps. (H) Stress-strain curve measurement of an IPN using amplitude sweep. (I) Measurement of Young’s modulus (N≥10), loss tangent (N≥10), stress relaxation (N≥2), linearity limit (N≥10), plasticity (N=3–7), and shear strengths (N≥10) of 7 distinct IPN formulations, where N indicates technical replicates. To refer to the different formulations, the following notations are used: ‘L’, ‘uL’, and ‘H’ represent LMW, unfiltered/unlyophilized LMW, and high-MW (HMW) alginate, respectively, and the number refers to the concentration of calcium crosslinker. For instance, ‘L3’ denotes an IPN formed with low-MW (LMW) alginate and 3 µM calcium crosslinking. (J) Measurement of Young’s modulus, loss tangent, linearity limit, and shear strengths of 5 distinct IPN formulations for primary T cell migration studies, with N≥5 technical replicates. (K) Heat map of coefficients of determination or R^2^ values from simple linear regression. (L) Measurement of Young’s modulus, shear strength, and loss tangent of human patient tissues, with N≥6 biological replicates from 7 patients. (I, J, L) *\*P* < 0.033, \*\**P* < 0.002, and \*\*\**P* < 0.001 by one-way ANOVA followed by Tukey’s post hoc analysis. All data are shown as means ± SEM.

Recent studies have generated deep insights into the physics of T cell migration in non-confining and partially confining microenvironments, but the mechanisms by which T cells generate migration paths in fully confining microenvironments has not been studied (*11, 12*). In non-confining microenvironments, such as microporous collagen gels, leukocytes are generally thought to mostly utilize an amoeboid or rounded mode of migration that does not rely heavily on adhesions, though they are guided by structural features of the matrix (*13-17*). In this mode, cells can utilize actin polymerization to expand into open spaces and contractility to squeeze their nucleus through any restricting matrix pores. However, when the pore size falls below 1–3 μm in diameter, this mode of migration becomes blocked (*18, 19*). T-cell migration in partially confining microenvironments has also been studied in contexts in which a migration path is pre-existing using microchannels, agarose gel overlays, where cells migrate at the interface between the agarose, or collagen gels that are microporous (*14, 18, 20-25*). In these contexts, T cells use the nucleus as a gauge to sense mechanical resistance. They harness topographical features to propel migration independent of adhesions, and generate lateral protrusive forces via polymerization of a central actin pool to “wedge” open the interface between two materials and protect the nucleus (*20-24*). How T cells generate migration paths in fully confining matrices with no pre-existing migration paths, however, is unknown. It is known that cancer cells, mesenchymal stem cells, and monocytes generate migration paths using either proteases, to biochemically degrade matrix, or mechanical forces, to mechanically open up channels, mediated by matrix mechanical plasticity (*26-29*). However, T cells are not thought to rely on proteases (*30*) and the nature of mechanical forces or matrix properties that might mediate migration path generation are unclear. Here we have asked how T cells generate migration paths in confining microenvironments, and what properties of the microenvironment ultimately govern and restrict migration (*12, 30*).

### Collagen-rich hydrogels with a range of mechanical properties

We first validated the nanoporosity of stromal ECM in normal bowel and pancreatic ductal adenocarcinoma (PDAC) tissues, representing healthy stroma and dense fibrotic tumor microenvironments, respectively. This was done by imaging glycosylated matrix components (e.g., proteoglycans) stained with Rhobo6, a recently developed glycan-binding dye (*5*), and collagen architecture via confocal fluorescence and reflectance microscopy, respectively (**Fig. 1B**). Analysis of visualized pores showed >98.5% of pores were below 1 µm in diameter in both healthy and diseased tissues, with a majority of the pore diameters below 400 nm in both cases, confirming that stromal matrices are predominantly nanoporous (**Fig. 1C, Fig. S1**). We note that this approach does not capture pores smaller than the ∼200 - 300 nm resolution limit of confocal microscopy, and therefore likely overestimates the actual pore size distribution.

Next, to investigate how ECM mechanical properties influence T cell migration through confining microenvironments, we developed interpenetrating network (IPN) hydrogel platforms that model the nanoporosity, mechanical properties, and collagen-rich characteristics of stromal ECMs in normal and fibrotic tissues (*31*). The platform consists of interpenetrating networks of type-I collagen and alginate. The collagen network mimics the microporous, fibrillar architecture of stromal collagen, while the alginate network enables tuning of overall IPN mechanical properties and roughly models other ECM components such as hyaluronic acid and proteoglycans. The collagen provides structural support and presents cell-adhesion ligands (e.g., α1β1, α2β1) for integrin-mediated T cell binding (*32*). In contrast, alginate is inert and lacks cell adhesion motifs, resists mammalian proteases, and forms a nanoporous mesh (*27, 31, 33, 34*). Given that the T cells are ∼11–13 μm in diameter, they must physically remodel the matrix and create micron-scale openings to migrate within the IPNs (*18*).

All IPNs used in this study were formed by blending type-1 collagen (1.5 mg/ml) with either high molecular weight (HMW) or low molecular weight (LMW) alginate (6.0 mg/ml), followed by crosslinking the alginate using calcium ions (**Fig. 1D**). Confocal microscopy revealed that the IPNs displayed a collagen fiber network with a micron-scale pore spacing (**Fig. 1E**). Confocal imaging of IPNs comprised of fluorescently labelled alginate indicated that the IPNs are nanoporous, with most pores ≤ ∼270 nm diameter (**Fig. 1F, Fig. S2**). This is consistent with previous work with similar IPN materials (*27, 31, 34*).

We used shear rheology to measure the range of material mechanical properties accessible in the IPN formulations. Young’s moduli and loss tangents were measured using oscillatory tests at small strains within the linear viscoelasticity limit and are indicative of stiffness and viscoelasticity respectively (**Fig. S3A**). Stress relaxation, another measure of viscoelasticity, was measured by applying a constant strain and quantifying the time for stress to be relaxed to half its initial value (*35*). Mechanical plasticity was characterized using creep and recovery tests, quantified as the ratio of irreversible strain after recovery to the maximum strain at the end of creep phase of the test (**Fig. S3B,C**) (*36*). Shear strength and linearity limit were measured using amplitude sweeps. In this method, a controlled sinusoidal strain was applied with incremental increases in amplitude, allowing the investigation of the material’s behavior from viscoelastic (non-destructive) to plastic and failing (destructive) deformation range (**Fig. 1G**). The linearity limit, defined as the strain at the upper limit of the linear relationship between stress and strain, was measured as the strain when G’ reached ±5% of its initial value. The shear strength represents the maximum oscillatory shear stress value at the point when the material begins to fail (**Fig. 1H**).

By systematically altering the alginate molecular weight (MW) and crosslinker concentration, we developed two sets of IPN hydrogels with a broad range of mechanical properties, including stiffness, viscoelasticity, plasticity, linearity limit, and shear strength that span a lower and higher range of shear strength, respectively. First, a set of seven distinct IPNs formed exhibited the following range of bulk mechanical properties: 500–4,000 Pa in Young’s modulus, 0.06–0.2 in loss tangent, 1–30 hours in stress relaxation half time, 0.03–0.13 in linearity limit, 0.09–0.25 in plasticity, and 50–600 Pa in shear strength (**Fig. 1I**). A second set of five distinct IPNs, which span a broader range of shear strengths, exhibited the following range of bulk mechanical properties: 1–5 kPa in Young’s modulus, 0.06–0.1 in loss tangent, 0.03–0.15 in linearity limit, and 0.3–7 kPa in shear strength (**Fig. 1J**). **Table S1** summarizes the compositions of these twelve IPNs. Notably, IPNs formed with HMW alginate exhibited shear strengths nearly an order of magnitude greater than their LMW counterparts at equivalent crosslinker concentrations. This is likely because the longer HMW chains create a more entangled and robust network that is more resistant to mechanical failure. Conversely, the shorter polymer lengths of LMW alginate result in a less interwoven network, making it more susceptible to tearing. Importantly, correlation analyses showed that no two parameters were coupled across these IPN sets (**Fig. 1K, Fig. S4**). Thus, each of these mechanical properties represent distinct behaviors of the material, and studies across matrices should allow determination of the independent correlation of each property to cell migration characteristics within the IPN.

To establish the physiological relevance of the range of shear strength accessible in this platform, since shear strength has not been studied previously, we measured the shear strength of of human tissues. Fresh normal bowel, normal pancreas, and PDAC tissues were analyzed using bulk rheology. These tissues displayed average properties spanning the following ranges from normal tissues to PDAC: Young’s modulus from 1 to 12 kPa; loss tangent from 0.2–0.4; and shear strength from 2 kPa to 60 kPa (**Fig. 1L, Fig. S5**). Thus, the mechanical profiles of the IPNs platform spans a portion of the range of stiffness and shear strength observed from compliant, normal bowel or pancreas tissues to stiff, fibrotic pancreatic cancer tissues.

### Matrix shear strength emerges as the strongest predictor of T cell migration

With these material platforms, we investigated how the matrix mechanical properties impact the 3D migration of T cells. Our study was conducted in two parts, first with Jurkat T cells and then with primary human T cells. Both Jurkat and primary T cells exhibited spontaneous random migration without requiring an externally applied chemokine gradient.

First, we examined Jurkat T cells using the first set of seven IPNs. The cells adeptly migrated through the nanoporous IPNs formed with LMW alginate and exhibited amoeboid-like morphology (**Movie S1)**. However, they were arrested and remained spherical in the IPNs with HMW alginate and higher levels of crosslinking (**Fig. 2A, Movies S2,3**). To quantify the motility, Jurkat T cells were fluorescently labeled and encapsulated in the seven distinct IPN formulations and tracked for a duration of six hours through time-lapse confocal microscopy (**Fig. 2B**). Several motility metrics were evaluated: the mean and maximum speeds of cell migration, track length (cumulative distance of all trajectory segments), track displacement length (straight-line start-to-end distance of the entire trajectory), and the fraction of motile cells (**Fig. 2C, Fig. S6**). Subsequent correlation analysis aimed to discern the influence of each hydrogel mechanical property on these motility descriptors (**Fig. 2D, Fig. S7**). Strikingly, among the mechanical parameters encompassing the entire range of material behaviors, shear strength exhibited the strongest correlation with every single metric of migration (**Figs. 2D,E, Fig. S7)**. Jurkat T cells became arrested in IPNs with shear strength above a threshold of around 500 Pa, indicating the limits of migration. In contrast, neither stiffness nor viscoelasticity showed a strong correlation with migration, indicating these properties do not regulate cell migration, at least across the range tested.

**Fig. 2.**
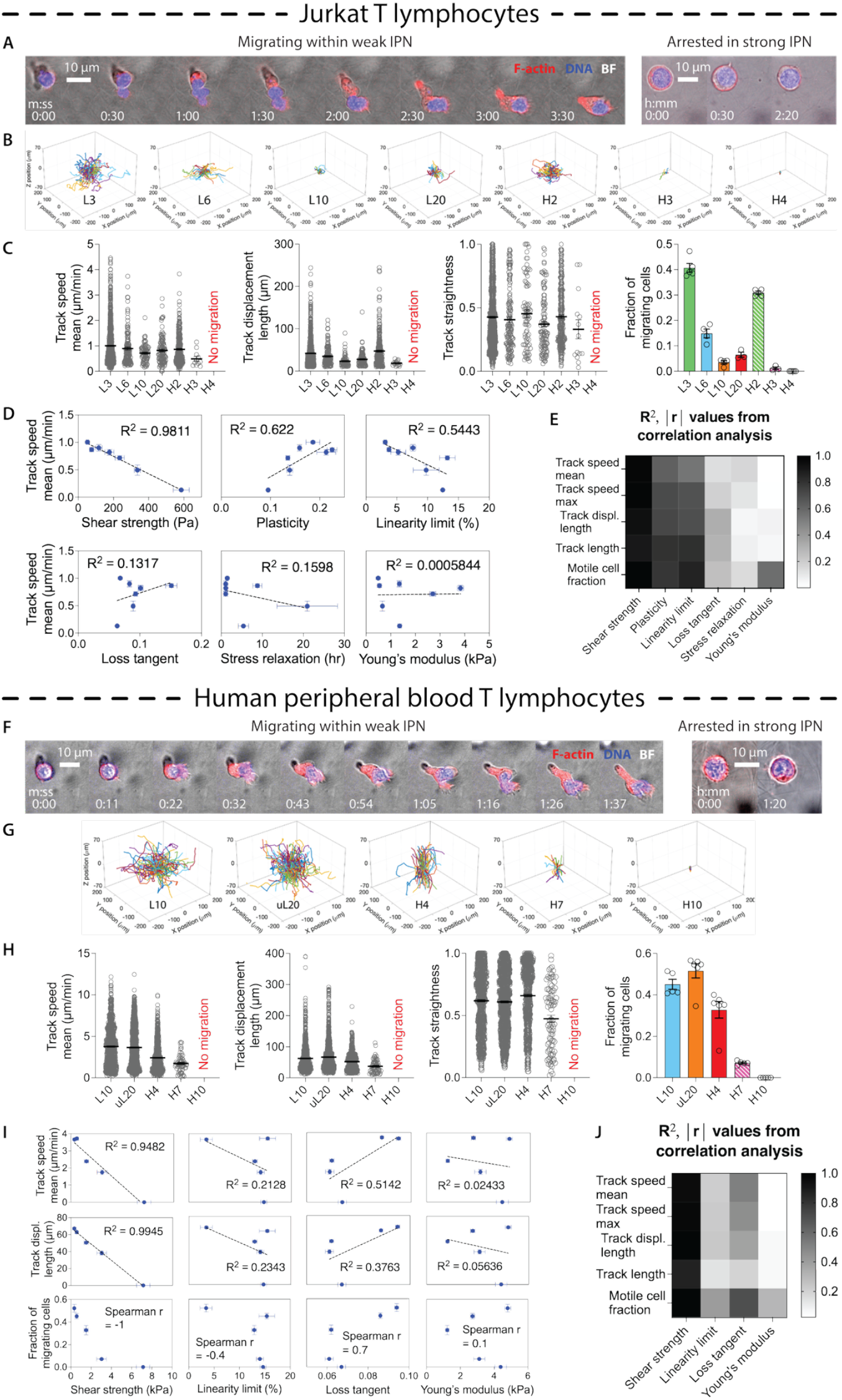
Matrix shear strength, not stiffness or viscoelasticity, correlates with T cell migration. (A) Time-series micrographs showing Jurkat T cells (left) migrating within a weak IPN and (right) arrested in a strong IPN. (B) Cell migration tracks observed in 7 distinct IPNs during a 6-hour assay. (C) Cell motility metrics measured per each IPN: track speed mean, track displacement length, track straightness, and fraction of migrating cells of all tracked cells. n≥281 for each condition; N≥3 replicates; from ≥2 independent experiments. (D) Simple linear regression applied to assess correlation of the cell motility and the IPN mechanical properties. R square values are shown on the graph. (E) Heatmap showcasing the correlation coefficient values derived from the analysis. (F) Time-series micrographs showing primary T cells (left) migrating within a weak IPN and (right) arrested in a strong IPN. (G) Cell migration tracks observed in 5 distinct IPNs during 3-hour duration assay. (H) Cell motility metrics measured per each IPN: track speed mean, track displacement length, track straightness, and fraction of migrating cells of all tracked cells. n≥1217 for each condition; N≥5 replicates. (I) Simple linear regression and Spearman’s rank correlation applied to assess correlation of the cell motility and the IPN mechanical properties. R square (linear regression) or r values (Spearman’s) are included on the graph. (J) Heatmap showcasing the correlation coefficient values derived from the analysis.

We then performed similar experiments with primary human T cells. We used activated human peripheral blood T cells, including ∼60% CD4+ and ∼40% CD8+ T cells (**Fig. S8**). Because activated primary T cells are much more powerful migrators—capable of migrating through hydrogels that arrested Jurkat cells—we utilized a second set of five IPNs spanning a much higher shear strength range (up to 7 kPa) to identify their mechanical migration threshold. Primary T cells stained with a fluorescent probe were encapsulated in the five distinct IPN formulations and tracked for a duration of three hours through time-lapse confocal microscopy. Primary T cells exhibited a similar trend of migration and morphologies to Jurkat T-cells: they adeptly migrated through IPNs with relatively low shear strengths, exhibiting amoeboid-like morphology, while they became arrested and remained spherical in the IPNs with high shear strengths (**Fig. 2F,G, Movies S4–7**). The motility characteristics of primary T cells were measured (**Fig. 2H, Fig. S9**). Again, matrix shear strength was most strongly correlated with the motility (**Fig. 2I,J, Fig. S10**). As with the Jurkat cells, stiffness and viscoelasticity did not correlate with migration. Despite the mixed CD4+ and CD8+ composition of the primary T cell population, we did not observe any bimodal distribution in the motility data, indicating that T cell motility in this context is likely subtype independent.

Taken together, these results demonstrate that matrix shear strength, rather than stiffness or viscoelasticity, is the key mechanical property governing the 3D migration of both Jurkat and primary T cells. These data indicate that the inability to induce material failure (i.e. fracture the matrix) ultimately limits T cell migration in confining ECMs.

### T cell migration depends on adhesions, actin polymerization and myosin contractility

We next sought to uncover the role of integrin-mediated adhesions and cytoskeletal machinery in driving T cell migration through confining matrices. Firstly, the role of adhesions was examined by comparing Jurkat cell motility in collagen-free alginate gels versus IPNs. We prepared two pairs of gels matched by shear strength: a ‘weak’ pair (∼100 Pa) and a ‘strong’ pair (∼250 Pa) (**Fig. 3A**).

**Fig. 3.**
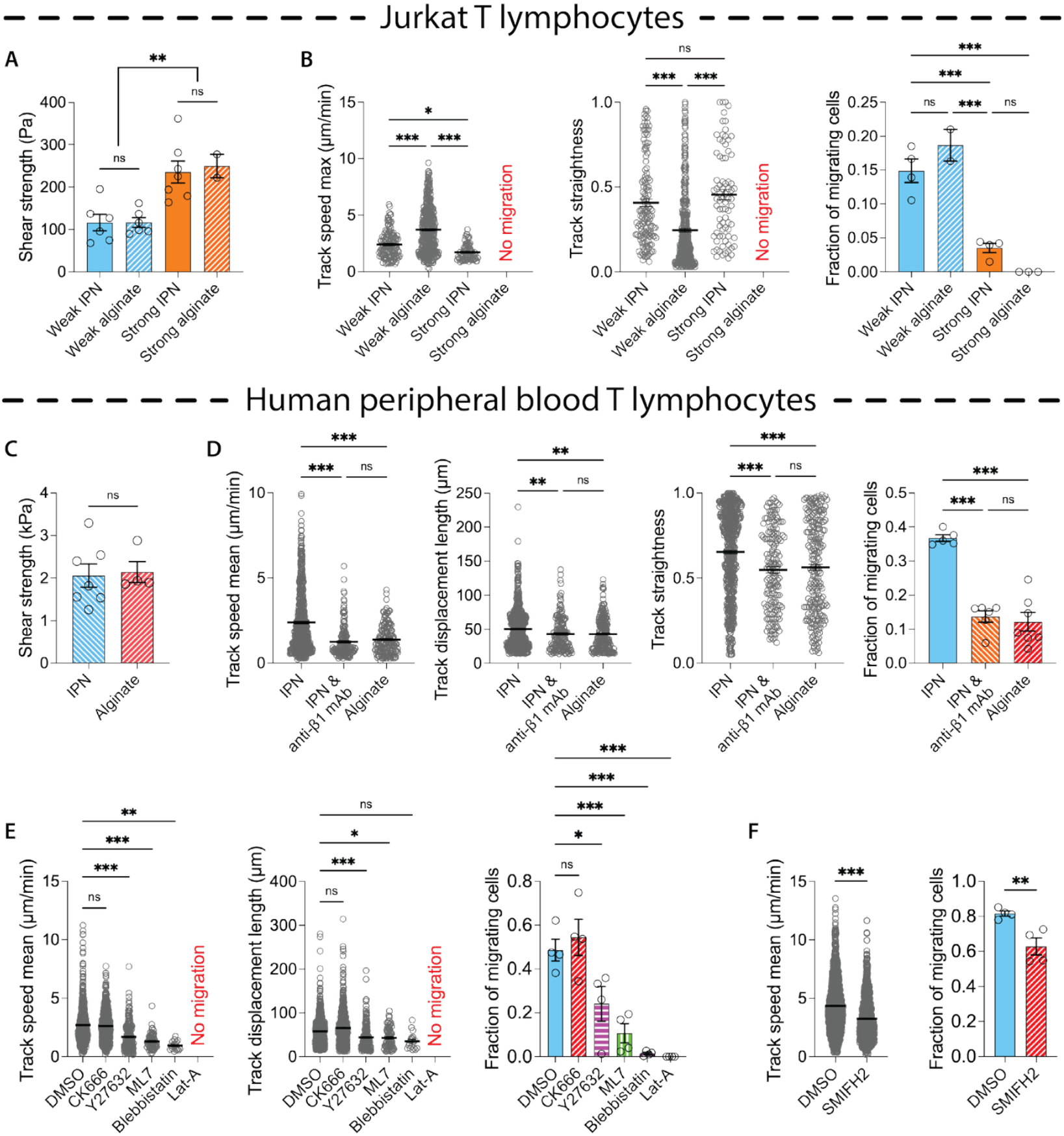
Roles of β1 integrin-mediated adhesions, actin polymerization, myosin, and formin activity in T cell migration. (A) Measurement of shear strengths of the two pairs of IPNs and alginate hydrogels; N≥4 technical replicates; Ordinary one-way ANOVA followed by Tukey’s post hoc analysis. (B) Cell motility metrics measured per each hydrogel: track speed max, track straightness, and fraction of migrating cells of all tracked cells. Total number of cell tracks n=1,070, 451, 2,167, 2,057 for weak IPN, alginate, strong IPN, alginate respectively. N≥2 replicates. (C) Measurement of shear strengths of the pair of IPN and alginate hydrogels; N≥4 technical replicates; Unpaired t test. (D) Cell motility metrics measured per each case (IPN, IPN & treated with *β*1 integrin monoclonal antibody, alginate): track speed mean, track displacement length, track straightness, and fraction of migrating cells. Total number of cell tracks n=1953, 1281, 2110 for IPN, IPN & anti-1 mAb, alginate respectively; 5-7 replicates; 2 experiments. (E) Cell motility metrics measured per each case of pharmaceutical inhibition: track speed mean, track displacement length, and fraction of migrating cells. Total number of cell tracks n=1536 (DMSO), 1114 (CK666), 817 (Y27632), 615 (ML7), 1260 (Bleb), 1260 (Lat-A); 4 replicates; 2 experiments. (F) Motility metrics measured for cells encapsulated within HMW IPNs treated with SMIFH2: track speed mean (unpaired, two-tailed Mann-Whitney U test) and fraction of migrating cells (unpaired t test); n=2078 for DMSO and 1619 for SMIFH2; 4 replicates; 2 experiments. (B,D,E) by One-way ANOVA followed by Tukey’s (fraction of migrating cells) and Kruskal-Wallis & corrected Dunn’s test (the other motility metrics) post hoc analysis. (A-F) All data are shown as means ± SEM; *\*P* < 0.033, \*\**P* < 0.002, and \*\*\**P* < 0.001.

The analyses reveal a dual, context-dependent role for adhesions. In the ‘strong’ gels, adhesions were critical for migration; Jurkat cells were completely immobilized in the pure alginate but achieved moderate, straighter migration in the IPN counterparts (**Fig. 3B**). This suggests that adhesive interactions are necessary for cells to generate sufficient traction to migrate in high-resistance environments. Conversely, in the ‘weak’ gels, adhesions appeared to interfere with efficient migration, as cells migrated slower in the IPNs than the adhesion-free alginate counterparts (**Fig. 3B**). This indicates that these same adhesive interactions may interfere with efficient amoeboid migration when the physical barrier is low.

We confirmed this principle with primary T cells using a pair of pure alginate and IPN gels matched by a shear strength of ∼2 kPa (**Fig. 3C**). The cells exhibited vastly superior motility—including a higher motile fraction, speed, travel distance, and directionality—in the IPNs compared to the alginate counterparts (**Fig. 3D**). Crucially, β1-integrin blockades in the IPNs reduced migration to the low levels observed in the alginate, confirming that T cells leverage adhesions to generate stronger forces necessary for efficient and linear path generation.

We next sought to identify the key cytoskeletal drivers of primary human T cell migration using a panel of pharmacological inhibitors. Depolymerizing filamentous actin (F-actin) with latrunculin A completely blocked cell migration, highlighting the essential role of F-actin in migration (**Fig. 3E**). Inhibition of myosin contractility, via ML7 or blebbistatin, and the Rho pathway, via the Rho kinase inhibitor Y27632, significantly diminished migration speed, travel distance, and the fraction of migratory cells, indicating the role of contractility (**Fig. 3E, Fig. S11A**). To further assess the role of actin polymerization, the actin nucleation promotion factors Arp2/3 complex and formins were inhibited via CK666 and SMIFH2 respectively. We observed that inhibition of the Arp2/3 complex, which generates branched actin networks, had no significant impact on cell migration (**Fig. 3F, Fig. S11A**). In contrast, inhibition of formins, which generate linear actin filaments, reduced migration speed, directionality, travel distance, and the fraction of migrating cells, implicating the role of formin-mediated actin polymerization (**Fig. 3F, Fig. S11B**). Cell viability was not significantly affected by any of the drug treatments (**Fig. S12**).

Together, these studies demonstrate that T cell migration through confining matrices relies on formin-driven actin polymerization and Rho-mediated actomyosin contractility to generate cellular forces for migration, and that T cells utilize β1-integrin mediated adhesions to generate greater migration forces in matrices with a higher shear strength.

### T cells rupture matrix via a breaststroke-like movement of actin-rich protrusions

Finally, we examined the hypothesis that T cells induce material failure in order to generate a migration path in confining collagen rich matrices by closely examining the dynamics of the migration process. In brightfield live-imaging, fingerlike protrusions were consistently observed forming at the leading edge. The protrusions moved in a divergent fashion at the leading edge and then moved towards the cell rear as they rounded around the leading edge and into the cell sides (**Fig. 4A, Movie S8**). Fluorescence live-imaging of labeled F-actin revealed that these fingerlike protrusions are actin-rich protrusions that extend into the hydrogel (**Fig. 4B, Movie S9**). Particle imaging velocimetry analysis of the actin dynamics measured the velocity of the divergent and then rearward actin movement originating from the leading edge of the migrating cells. The accumulation of actin moving retrograde resulted in a dense actin punctum at the trailing edge (**Fig. 4C**). Notably, the maximum retrograde actin flow velocity (∼4 µm/min) essentially matches the cell’s translocation speed (Fig. 2H) suggesting a high efficiency motion. This motion of finger-like protrusions was reminiscent of the breaststroke in swimming and is hereafter referred to as a breaststroke-like motion. Note that T cells were previously shown to undergo a breaststroke-like motion during swimming in fluid (*37*).

**Fig. 4.**
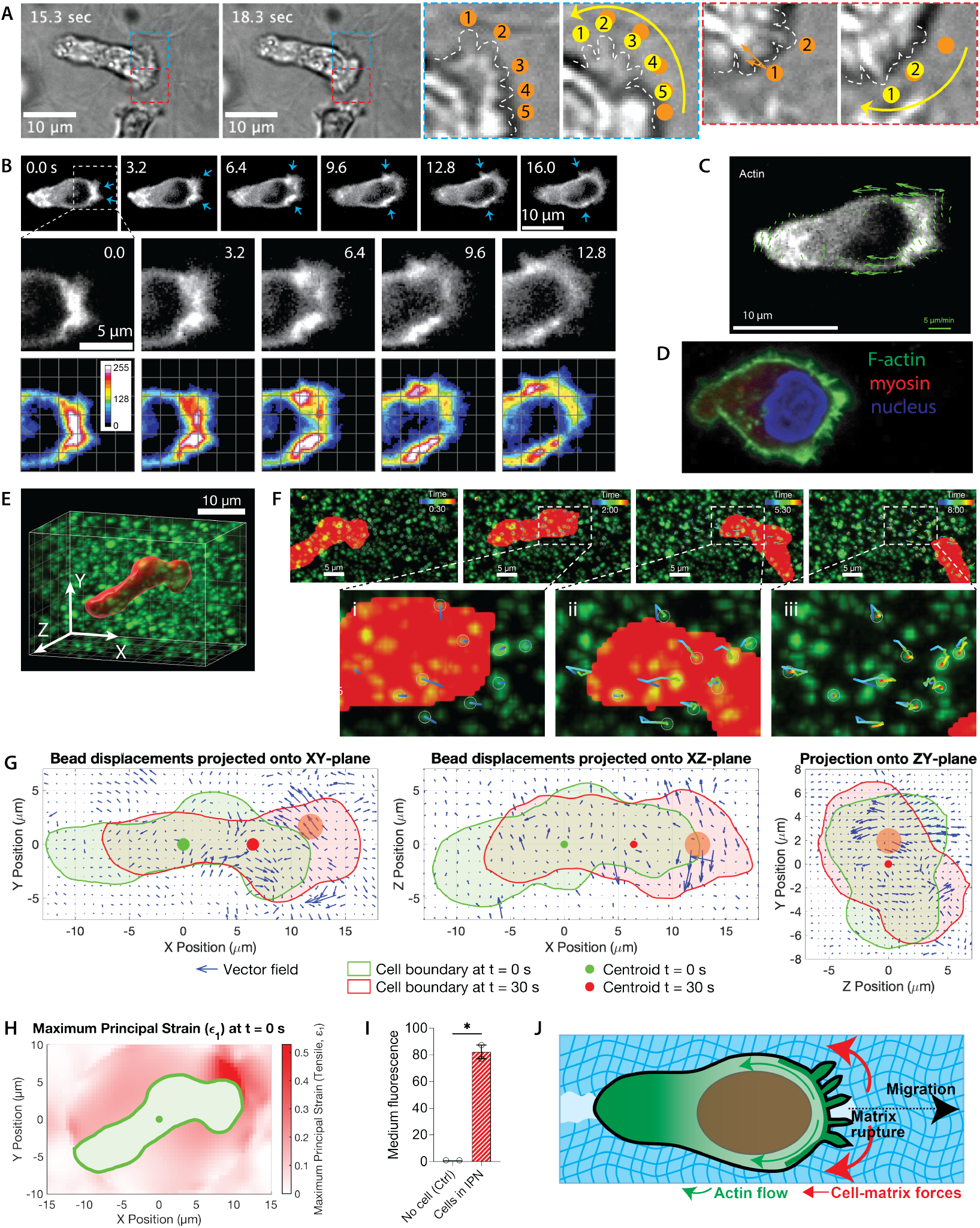
T cells tear apart matrix via a divergent, breaststroke-like movement of actin-rich protrusions at the leading edge. (A) Brightfield images showing the coordinated, breaststroke-like movement of finger-like protrusions at the leading edge of a T cell migrating in an IPN. Zoomed-in frames (outlined in green and red) highlight the numbered protrusions displaced between 15.3 and 18.3 seconds. (B) Time-lapse fluorescence micrographs showing F-actin dynamics in a migrating T cell; the bottom row visualizes bright F-actin clusters using a colormap of fluorescence intensity. (C) F-actin movement speed quantified by particle image velocimetry. (D) Immunofluorescence micrograph of a migrating T cell. (E) 3D Imaris reconstruction of a T cell in an IPN embedded with fluorescent beads. (F) Time series snapshots tracking bead displacements induced by T cell migration in Imaris, with insets depicting representative tracks over (i) 30 seconds, (ii) 4 minutes, and (iii) 6 minutes 30 seconds. (G) Vector maps of 3D bead displacements over the first 30 seconds, projected onto the XY, XZ, and YZ planes, alongside cell centroids and boundaries. The orange circle indicates the estimated location of cell’s leading edge, identified by the convergence point of the displacement vectors and its proximity to the region of maximum principal strain. The data shown are from a representative time point for one of four cells analyzed. (H) 3D maximum principal strain measurements mapped over the XY plane. (I) Alginate fluorescence in media for cell-encapsulating IPNs versus cell-free IPNs. The fluorescence intensity values were normalized by those of cell-free samples (control). n=2 from 2 experiments. (J) Proposed migration model. Cells create actin-rich protrusions at the leading edge which undergo a breaststroke-like motion, locally rupturing matrix to open a path to migrate. Green arrows: F-actin movement. Red arrows: cell-generated matrix forces. Data are shown as means ± SEM; *\*P* < 0.033, \*\**P* < 0.002, and \*\*\**P* < 0.001 from paired two-tailed t test.

To visualize the actin and myosin architecture driving this migration, we performed immunofluorescence staining on fixed migrating cells. This analysis revealed the detailed structure of the actin-rich protrusions around the cell boundary as well as the diffuse, uniform distribution of myosin throughout the cytoplasm.

We next investigated whether the breaststroke-like motion could rupture the ECM, thereby generating a migration path. To study this, we embedded fluorescent beads into the hydrogel to monitor hydrogel deformations around migrating T cells (**Fig. 4E**). Individual beads were tracked via time-lapse confocal microscopy and Imaris analyses, revealing a complex pattern of movement (**Fig. 4F, Movie S10**). Initially, beads near the cell’s leading edge were pulled backward and outward in a divergent manner, a motion consistent with the cell’s breaststroke-like action (**Fig. 4F-i**). Subsequently, as the cell advanced through the newly created pathway, the beads were dragged forward by the cell’s movement (**Fig. 4F-ii**). Finally, as the cell migrated past and away from the bead, the beads were displaced backward toward its original vicinity (**Fig. 4F-iii**). This final movement likely reflects the partial recovery of the hydrogel, which involves elastic recoil and tunnel constriction upon the removal of tension and compression respectively.

To quantify these observations and reveal more precisely the nature of matrix deformation associated with migration, 3D hydrogel deformation and traction strain maps were generated from the bead displacements (**Fig. 4G, Fig. S13**). These maps revealed two distinct patterns. At the cell’s leading edge, the matrix strain exhibited a radially divergent, rearward pattern, consistent with the cell’s “breaststroke” motion. In contrast, beads near the cell’s centroid and rear were displaced forward, indicating a forward strain of the matrix. This movement is likely caused by shear stress from the advancing cell, consistent with the observations in Fig. 4F and Movie S12. Importantly, the principal strains in the matrix were greatest at cell’s leading edge, estimated to be on the order of 0.3–0.5, a range well within the threshold expected to induce matrix rupture (**Fig. 4H**).

To determine if migrating cells leave behind permanent tunnels, we imaged them in fluorescent-alginate IPNs. Interestingly, the fluorescence in the path recovered immediately after the cell passed, making the tunnels appear transient (**Movies S11,12**). We hypothesized this was caused by mechanically liberated fluorescent alginate molecules diffusing into the medium and filling the void. This hypothesis was confirmed by finding that supernatant media from cell-laden hydrogels were ∼80-fold more fluorescent than from acellular controls (**Fig. 4I**), while the bulk gel fluorescence intensity decreases over time (**Movie S12**). This demonstrates that migrating T cells mechanically rupture the matrix, and the liberated polymer debris diffuse into the media and the newly formed channel.

From these data, we infer that the breaststroke like motion of the actin rich protrusions generates divergent mechanical stresses on the matrix that causes matrix rupture, thereby generating a path for the cell to migrate into (**Fig. 4J**).

## Discussion

Our work identifies matrix shear strength as a key mechanical property that regulates T cell migration through confining microenvironments. While the field of mechanobiology has extensively explored the roles of matrix stiffness and degradability on 3D cell migration, and more recently viscoelasticity and mechanical plasticity, our results indicate that a matrix’s resistance to rupture—its shear strength—presents the critical barrier that arrests cell movement when it exceeds a cell’s force-generating capacity (*11, 12*). This finding establishes a fundamental biophysical principle on cell-matrix interactions.

To overcome this physical barrier, we found that T cells utilize a previously undescribed migration mechanism, which we term a “breaststroke”, to make their way through highly confining microenvironments In this process, cells extend actin-rich, finger-like protrusions and then use a divergent stroke at the leading edge to physically tear apart the matrix to generate a migration path. This mechanism is distinct from other migration modes previously described (*13-18, 20-23*). These include exocrine gland resident CD8+ T cell migration led by blebs at the leading edge (*25*), or migration in less-confining contexts that involves polymerization of a central F-actin pool to push laterally (*23, 24*). It also differs from protease-independent mechanisms used by other cell types, such as the repeated extension-retraction cycles of invadopodia in cancer cells (*27*), the nuclear piston mechanism of mesenchymal stem cells (*28*), or the primarily protrusive forces generated at the leading edge by monocytes (*29*). Remarkably, T cells migrate approximately an order of a magnitude faster than these other cells in confining hydrogels, suggesting that this breaststroke like motion used by T cells may be much more effective at clearing matrix.

The high efficiency of this migration mechanism, with the cell’s translocation speed nearly matching the maximum speed of its protrusion strokes, suggests strong evolutionary pressure. This nearly 100% efficiency in migrating through ECMs shows sharp contrast to the known inefficiency of T cell swimming in fluid. In fluid, the ratio of cell speed to the membrane protein speed (i.e., stroke speed) is only 20%, far lower than the ∼67% efficiency reported for single-celled organisms driven by cilia or membrane treadmilling (*37*). Aoun et al. attributes this low efficiency to only a small fraction of the T cell membrane actively participating in stroke generation (*37*). However, when migrating through a confining ECM, a T cell requires only sufficient protrusions to “grab” matrix fibers, allowing stroke speed to translate directly into rapid forward movement. We propose that T cells evolved this powerful breaststroke primarily for efficient navigation of soft tissue ECMs, while retaining sufficient capability to swim in fluidic environments (e.g., amniotic or ocular fluid), enabling them to patrol virtually every solid and fluid compartment in the body.

Although T cells often migrate along preexisting paths in vivo, fibrotic tumor microenvironments likely require them to penetrate highly confining ECMs using this migration mode. Indeed, our analysis of matrix pore sizes in human pancreatic adenocarcinoma (PDAC) tissues point to pore structures that are predominantly submicron-sized or highly confining. Interestingly, we found that the shear strength of these tissues was more than 20 times higher than those of normal pancreas tissues (**Fig. 1I**). Furthermore, primary T cells were immobile in IPNs exhibiting the high shear strength of PDAC but mobile in those exhibiting the low shear strength of normal pancreatic tissues (**Fig. 2H**). These results suggest that the limited ability of T cells to infiltrate the PDAC microenvironment may be in part because the changes in shear strength occurring during pancreatic cancer progression could limit the physical ability of the T cells to infiltrate these matrices.

The implications of these findings are broad. First, they suggest that shear strength should be considered a critical parameter in the design and characterization of biomaterials for tissue engineering, regenerative medicine, and immunotherapy, as it may be a more accurate predictor of cell infiltration than stiffness or pore size alone. Second, the ability to overcome a matrix’s shear strength may be a relevant factor in the migration of other cell types, including cancer cells, fibroblasts, and other immune cells, and this warrants further investigation. More broadly, our study suggests that shear strength deserves consideration as a key variable in mechanobiology, potentially playing a role in diverse dynamic cell-matrix interactions, including cell spreading, protrusion extension, and morphogenesis.

## Acknowledgments

We thank Dr. Jake Song for providing fluorescently labeled alginate, and Dr. Kayvon Pedram, Janelia Research Campus, Howard Hughes Medical Institute (HHMI), for gift of Rhobo6 dye and guidance on its use.

## Funding

National Science Foundation Postdoctoral Fellowship in Biology 2209411 (BH)

Stanford Cancer Institute Postdoctoral Fellowship Award (BH)

National Institutes of Health grant T32 5T32DK007056-49 (NJT, BH)

National Institutes of Health grant R01 CA277710 (NJT)

National Institutes of Health grant R37 CA214136 (OC)

National Institutes of Health grant R01 CA290021 (OC)

National Institutes of Health grant R01 GM14853 (OC)

## Author contributions

Conceptualization: BH, OC

Methodology: BH, PX, BJ, CA, MK

Investigation: BH, PX, BJ, CA, MK

Funding acquisition: BH, NT, OC

Project administration: DD, PB, NT, OC

Supervision: DD, PB, NT, OC

Writing – original draft: BH, OC

Writing – review & editing: All authors

## Competing interests

The authors declare no competing interests.

## Data, code, and materials availability

All processed data are included in supplementary information. Raw data is available upon request.

## Supplementary Materials

Materials and Methods

Supplementary Text

Figs. S1 to S13

Table S1

Movies S1 to S12

## Notes

### Competing Interest Statement

The authors have declared no competing interest.

### Summary of Updates

Additional main and supplementary data and Methods & Materials descriptions have been added. The manuscript has been updated accordingly.

