## Supplementary Material for "T Cells Tear Apart Confining Extracellular Matrix Via a Breaststroke-like Motion to Generate Migration Paths"

**The PDF file includes:**

Materials and Methods
Figs. S1 to S13
Tables S1

**Other Supplementary Materials for this manuscript include the following:**

Movies S1 to S12

### Materials and Methods

**Cell culture.** All cells were grown and maintained at 37°C in a standard humidified incubator with 5% $CO_2$ atmosphere and routinely tested for mycoplasma contamination.

**Jurkat T cell culture and reagents**. Jurkat T cells were grown and maintained in suspension culture in RPMI 1640 medium containing 2 mM L-glutamine and 110 mg/L sodium pyruvate (Gibco, A1049101) supplemented with 10% fetal bovine serum (Hyclone), 1% Penicillin/Streptomycin (Life Technologies), and 100 μM 2-mercaptoethanol. Cells used in this study were between passages 8 and 10. Cells were maintained at sub-confluency and passaged every two days. For cryopreservation, cells were pelleted by centrifugation, resuspended in freezing medium consisting of 90% FBS and 10% dimethyl sulfoxide (DMSO; Tocris Bioscience), and placed in cell-freezing containers at −80 °C overnight prior to transfer to liquid nitrogen for long-term storage.

**Primary T cell culture and reagents**. Primary human T cells were purchased from STEMCELL Technologies (#70024). According to the vendor, the cells were isolated from peripheral blood mononuclear cells (PBMCs) using negative immunomagnetic separation and were mostly naïve T cells with ~60% CD4+ and ~40% CD8+ subtypes. Upon receipt, cells were initially expanded in culture for 12 days and cryopreserved in multiple aliquots for subsequent use. T cell activation was induced by supplementing cultures with 2–10 ng/mL human recombinant interleukin-2 (IL-2; STEMCELL Technologies, #78036) and 25 μL/mL anti-CD3/CD28 antibody cocktail (STEMCELL Technologies, #10971). Cells were grown and maintained in suspension in RPMI 1640 medium (Gibco, A1049101) supplemented with 10% fetal bovine serum (Hyclone), and 1% Penicillin/Streptomycin (Life Technologies), 100 μM 2-mercaptoethanol, 1× non-essential amino acids (Gibco, #11140035) and 50 μM ascorbic acid (Sigma-Aldrich, A8960). In this study, we only used the cells thawed from the first expanded aliquots of passage 1 and the cells used in this study were between passages 2 and 3. In all migration assays where motility was assessed, cells were activated two days before using the IL-2 and anti-CD3/CD28 activator.

**Alginate preparation.** Low molecular weight (LMW) ultra-pure sodium alginate (<75 kDa; Provona UP VLVG, NovaMatrix) and high-MW alginate (280 kDa; Protanal LF 20/40, FMC Biopolymer) were used. Stock solutions were prepared by dissolving each alginate in deionized water, followed by sterile filtration and lyophilization. The resulting dry powder was then reconstituted in serum-free Dulbecco's modified Eagle's medium (DMEM; Gibco) to a concentration of 3.5 wt% (35 mg/mL). To formulate an IPN with stiffness comparable to PDAC and H10 IPNs but lower shear strength, unfiltered and unlyophilized LMW (designated "uLMW") alginate was directly dissolved from the stock bottle in DMEM. Based on the altered mechanical properties of uL20 IPNs made with this uLMW compared to filtered, lyophilized alginate, we assume that this modified protocol retained a greater proportion of higher molecular weight chains. Alginate hydrogels were prepared using either LMW, uLMW, or HMW formulations at a final concentration of 0.6 wt% (6 mg/mL). For preparation of fluorescent alginates, fluorescein amine isomer

(Acros Organics) was conjugated to alginate using standard carbodiimide chemistry at 37.74 μM (38). A full description of all the formulations used is included as Supplementary Table 1 (**Table S1**).

**Hydrogel formation and cell encapsulation.** For alginate and IPN gels, alginate (reconstituted in DMEM solution as described above) was delivered to a 1.5 mL Eppendorf tube at room temperature (21°C). For collagen and IPN gels, rat tail collagen I (Corning), was neutralized with 10X DMEM to have pH of ~7 at 4°C. Neutralized collagen was added to the alginate and carefully mixed >30 times with a pipette to avoid generating bubbles. Extra DMEM was added to ensure 6.0 mg/ml - 1.5 mg/ml alginate-collagen final gel concentration. For 3D migration assays, cells were added to the hydrogel solutions. The mixture of the polymers and/or cells was transferred to a separate 1 mL Luer lock syringe (polymers syringe).

Next, calcium sulfate slurries in DMEM were added to a 1 mL Luer lock syringe (Cole-Parmer). The calcium sulfate solution was shaken to mix the calcium sulfate evenly, and it was then coupled to the polymers syringe with a female-female Luer lock (Cole-Parmer), taking care not to introduce bubbles or air in the mixture. Finally, the two solutions were rapidly mixed together with 2-12 pumps on the syringe handles and instantly deposited into a well in a 6-well glass-bottom dish (Thermo Scientific). Gelation of all hydrogels was completed at 21ºC for 30 min before media was added to the wells.

Cells were resuspended in growth medium containing CellTracker cytoplasm fluorophore (ThermoFisher), centrifuged, and subsequently suspended in DMEM. Cell concentrations were quantified using a Vi-Cell Coulter counter (Beckman Coulter). For migration and inhibitor assays, cells were seeded at $\sim 2 \times 10^6$ cells/mL. For mechanistic studies, including analyses of actin dynamics and bead displacement, cells were seeded at $\sim 1 \times 10^7$ cells/mL to facilitate high-magnification imaging and ensure sufficient numbers of migrating cells within the field of view.

**Extraction of human tissues from PDAC resection.** Fresh tissue samples were obtained from a PDAC tumor resection on the day of Whipple surgery. Both tumor and adjacent healthy bowel and pancreas tissues were identified and isolated by a pathologist. The bowel tissue was cut longitudinally to lay flat, and the epithelial layer on the luminal surface was carefully removed to expose the submucosa.

**Pore size measurement of tissues.** The tissues were sectioned into 2 mm-thick slices, from which 6 mm-diameter pucks were punched out for staining. Samples were maintained in DMEM supplemented with 10% FBS to preserve viability. Tissue pucks were first stained with Hoechst. The staining medium was prepared by diluting Hoechst (1:2000) in DMEM. The samples were immersed in this medium for 30 min at 37 °C and 5% $CO_2$ in the dark. After incubation, the medium was removed and samples were washed twice with PBS. Next, Rhobo6 staining was performed by immersing the samples in DMEM containing Rhobo6 (1:200). The samples were immersed for 1 h at 37 °C and 5% $CO_2$ in the dark. Because Rhobo6 is a reversible dye, imaging was performed immediately after staining without changing the medium.

For measuring the pore size of the IPNs, an IPN of 6 mg/mL fluorescent HMW alginate and 1.5 mg/mL type-1 collagen with 4 μM calcium ion crosslinker (i.e., H4 IPN using fluorescent alginate; total gel volume 400–450 μL) was prepared in a 6-well glass-bottom dish and hydrated with 2 mL of DMEM. After 10 h of equilibration, 500 uL of the supernatant medium—potentially containing fluorescent alginate molecules leached from the hydrogel—was removed and transferred to a 6-well glass-bottom dish for background fluorescence measurement.

All samples were imaged using a Leica SP8 laser-scanning confocal microscope in an environmental control chamber maintained at 37°C and 5% $CO_2$. A 25x 0.95-NA water-immersion objective was used for all images. All tissue samples were imaged within a few hours after staining to limit any effect due to cell death. IPN and background media images were acquired under identical imaging conditions (e.g., pinhole size, laser power, detector gain). For reflectance imaging, a 488 nm laser was used to visualize the overall ECM architecture.

Each image was converted into a binarized image by masking empty spaces and applying a threshold based on background intensity. The threshold was defined as the mean intensity of an empty peripheral region lacking cells or tissue, representing background fluorescence from Rhobo6 and medium autofluorescence. Pixels below this threshold were defined as "empty." The image was masked to include only tissue regions and then analyzed using a MATLAB program to measure pore sizes. The binary image was inverted so that pores corresponded to pixel value 0 and tissue to 1. Connected pore regions were identified, and the number of pixels per region was counted. Pixel counts were converted to physical areas using a pixel-to-area ratio of 1 pixel = 0.0516 $\mu m^2$ (as determined by magnification and imaging parameters). Assuming circular geometry, pore diameters were estimated from the measured areas. Both the histogram and cumulative distribution function (CDF) of pore sizes were plotted.

The IPN image (**Fig. S2A**) was binarized by applying a threshold value above the maximum background intensity (**Fig. S2B,C**). Pores were segmented and analyzed in Imaris to measure area and count (**Fig. S2D,E**).

**Mechanical characterization of hydrogels.** Hydrogels (IPNs, alginate, and collagen gels) were mechanically characterized using an DHR-2 stress-controlled rheometer (TA Instruments) with 25 mm parallel plate geometry. To prevent interfacial slip, both plates were coated with 0.1 mg/mL poly-L-lysine aqueous solution (10 min incubation followed by air drying) prior to the deposition of hydrogel solution. All rheological measurements were performed 21°C, maintained by a Peltier system to match the conditions used for cell encapsulation.

Alginate-bearing hydrogel precursor solutions were mixed with crosslinkers using dual-barrel syringes connected via Luer-Lock to ensure homogeneous blending. For collagen gels, 400 µL neutralized collagen was prepared. Approximately 400 µL of the hydrogel solution was deposited onto the lower PLL-coated plate. The upper plate was immediately lowered to achieve a 550 µm gap, forming a hydrogel disk between the two plates. Mineral oil (Sigma) was applied around the sample edges to prevent dehydration.

During gelation, the storage (G') and loss (G") moduli were monitored for 1 hour with continuous oscillations at a strain of 0.01 and frequency of 1 rad/s (**Fig. S1A**). The moduli recorded at the end of this period were used for all the analyses. For the IPNs used in Jurkat cell studies, measurements were performed at 1 rad/s. To ensure measurements were within the linear viscoelastic (LVE) regime, amplitude sweeps were performed to determine the upper linearity limit for each hydrogel type. The applied 1% strain was well below these limits for all the hydrogels used in this study. The complex modulus ($G^*$) was calculated as,

$$G^* = \left(G'^2 + G''^2\right)^{1/2}. \tag{1}$$

The elastic (Young's) modulus was then determined assuming a Poisson's ratio ($\nu$) of 0.5 using the equation,

$$E = 2(1 + \nu)G^*. \quad (2)$$

For stress relaxation measurement, a strain-creep test was conducted following this oscillatory test. A constant shear strain of 0.02 was applied for 2–20 hours, or until the measured stress decayed to half its initial value. All strains applied in these tests were confirmed to be below the LVE limits.

For plasticity measurement, creep and recovery tests were performed following the oscillatory test (**Fig. S1B**). A constant shear stress of 50 Pa (exceeding the yield strength) was applied for 30 minutes while recording strain over time (**Fig. S1C**). The sample was then unloaded (0 Pa), and strain recovery was monitored for 3 hours. Plasticity was quantified as the ratio of irreversible strain after recovery to the maximum strain at the end of the creep phase, or

$$plasticity = \varepsilon_{irreversible}/\varepsilon_{maximum}, \quad (3)$$

where $\varepsilon_{maximum}$ is the maximum strain at the end of the creep test, and $\varepsilon_{irreversible}$ is the residual strain after the recovery test.

For linearity limit and shear strength measurement, amplitude sweeps were performed following the oscillatory test. In this method, controlled sinusoidal strain at 1 rad/s frequency was applied with incremental increases in amplitude, allowing the investigation of the material's behavior from a non-destructive to a destructive deformation range. In general, when examining an unknown sample by an oscillatory test, an amplitude sweep must first be carried out in order to determine the linearity limit. Establishing these parameters is essential: all linear elastic and viscoelastic tests must be conducted within the LVE region, while plasticity measurements required applying stresses exceeding the yield strength.

The linearity limit is defined as the strain at ±5% deviation of plateau G' value. The shear strength represents the maximum oscillatory shear stress value at the point when the material begins to fail. Note the range of 9–25% in plasticity indicates that the IPNs recovered 75-91% strain after the removal of creep stress, while linear viscoelasticity and shear strength correspond to 100% and 0% strain recovery, respectively.

**Mechanical characterization of tissues.** For rheological measurements, tissues were sectioned into 1–2 mm thick slices, from which 8 mm diameter discs ("pucks") were punched. All tests were performed on an DHR-2 stress-controlled rheometer (TA Instruments) with an 8 mm parallel plate geometry at 21°C. For low-strain oscillatory tests to measure G' and G", samples were mounted without adhesive on plates covered with double-sided tape and Kimwipes. Slippage was prevented by a combination of friction (enhanced by an axial force of ~0.03 N) and water-mediated adhesion. For these tests, G' and G" were measured by applying 1% strain at 1 rad/s. For high-strain amplitude sweeps (up to 2000%) to measure shear strength, tissue pucks were affixed to the Kimwipe-covered plates using superglue (Gorilla). To prevent artifacts, a minimal amount of glue was applied and allowed to pre-thicken for 2–3 minutes via exposure to ambient moisture before the puck was mounted; this limited glue penetration and artificial hardening of the tissue. Care was also taken to prevent glue bridging between the top and bottom plates, either through the tissue or around its outer surface. To validate this adhesive method, the initial low-strain G' and loss tangent from the amplitude sweep were compared with the values from the non-adhesive oscillatory tests to confirm that no significant tissue hardening had occurred.

**Imaging cell migration.** All microscope imaging was done with a laser scanning confocal microscope (Leica SP8) and disk-spinning confocal microscope (Nikon) fitted with temperature/incubator control, suitable for live-imaging (37 °C, 5% $CO_2$). In live-cell time-lapse imaging used for motility measurements, CellTracker-labeled cells were tracked with a 10x NA 0.75 air objective for 3 hours (primary T cells) and 6 hours (Jurkat T). Additionally, 64 μm z-stack images were acquired every 2-5 minutes and imaging parameters were adjusted to minimize photobleaching and avoid cell death. For brightfield imaging that captured breaststroke-like motion of protrusions, primary pan-T cells were imaged every 100 ms with a Nikon 40×, NA 1.15 oil-immersion objective. For actin dynamics imaging, primary pan-T cells were labeled with SPY650-FastAct (Cytoskeleton, Inc., SC505; F-actin stain) at a 1:1,000 dilution for 4 h. Time-lapse images were acquired every 300–700 ms with a Nikon 40×, NA 1.15 oil-immersion objective. Heatmaps were produced by performing a maximum-intensity z-projection, applying a smoothing algorithm, and assigning colors using the 16-color lookup table in ImageJ.

**Confocal reflectance microscopy for collagen fiber characterization.** Alginate–collagen matrices were prepared in 6-well glass-bottom dishes and imaged using a laser scanning confocal microscope (Leica SP8) equipped with a 25×, NA 0.95 water-immersion objective. Samples were excited with a 488-nm laser, and reflected light was collected in reflectance mode.

**Imaris cell tracking algorithm.** For migration studies, the centroids of fluorescently labeled cells were tracked using the surfaces and spots functionalities in Imaris (Bitplane). Poorly segmented artifacts and cell debris were excluded from the analysis and drift correction was implemented where appropriate. A custom MATLAB script was used to reconstruct cell migration trajectory. Cells were considered arrested if they exhibited minimal motility over the measurement period (i.e., 3 hours for activated primary T cells and 6 hours for Jurkat T). Quantitatively, a cell was considered arrested if its track displacement length was below a specific threshold. This threshold was established for each experiment and ranged from 8–20 μm, depending on the effectiveness of the computational drift correction.

In certain HMW IPNs, which exhibit relatively high shear strength and permit only limited T cell migration, cells occasionally encountered dead-ends after generating a migratory path. Upon reaching such dead-ends, T cells reversed direction and retreated through the tunnels they had previously created, traveling at approximately twice their forward speed (**Movie S6**). Some T cells became confined within such tunnels and exhibited repeated back-and-forth movement.

Our objective was to measure motility associated with *de novo* path generation, not the movement of cells squeezing through pre-existing paths. Therefore, to ensure data accuracy, we excluded measurements associated with faster movement through pre-existing paths. First, we initiated cell tracking as soon as gel encapsulation, mostly within 1–2 hours, and utilized the Imaris user interface to manually review individual cell tracks. Tracks appearing to migrate through pre-existing tunnels or exhibiting back-and-forth movement were identified and excluded from the analysis. These exclusions were applied only to primary cells in the H4 and H7 IPNs and to Jurkat cells in the H3 IPN, where such behaviors were prominent in high shear strength gels that restrict migration. In contrast, in low shear strength gels using LMW alginate or H2 with low crosslinking, cells rarely encountered dead ends and typically migrated without reversing direction, so any such events would have negligible impact on the averaged motility measurements.

**Inhibition studies.** For all pharmacological inhibition experiments, inhibitors were added to the culture medium 1 h prior to the start of cell migration tracking. The following inhibitors were used: CK666 (5 µM; Sigma-Aldrich SML0006; Arp2/3 inhibitor), Y27632 (10 µM; STEMCELL 72304; ROCK inhibitor), ML7 (10 µM; Tocris Bioscience 4310, myosin light chain kinase inhibitor), Blebbistatin (10 µM; Abcam ab120425; myosin II inhibitor), Latrunculin A (200 nM; Tocris Bioscience 3973; F-actin polymerization inhibitor), and SMIFH (20 µM; Sigma-Aldrich S4826; formin inhibitor). ML7 was dissolved in ethanol, while all other inhibitors were prepared in dimethyl sulfoxide (DMSO; Sigma-Aldrich D8418). Stock solutions were diluted to their final concentrations in growth medium before being added to cell-encapsulating gels. Control samples received DMSO diluted 1:1,000 (v/v) in growth medium. Cell migration was then imaged using time-lapse confocal microscopy every 2–3 minutes for 3 h.

**Live/dead studies.** Cell viability was assessed using the LIVE/DEAD Viability/Cytotoxicity Kit (Invitrogen L3224). Hydrogels containing cells were incubated with 2 µM calcein-AM and 4 µM ethidium homodimer-1 for 45 min, followed by immediate imaging. Live cells exhibited green fluorescence due to esterase activity, whereas dead cells displayed red fluorescence indicating loss of plasma membrane integrity.

**Actin flow analysis in migrating cell using Particle Image Velocimetry.** Actin flow velocity fields were computed from sequential images at 3.2 second intervals using PIVlab in MATLAB (39). To preprocess the image sets, a high-pass filter and contrast limited adaptive histogram equalization (CLAHE) were applied, and regions outside the cell body were manually masked. A multi-pass FFT window deformation algorithm with a 32x32 pixel interrogation area was used to track actin movement. Velocity vector fields were visualized using a custom MATLAB script.

**Bead displacement assays for measuring cell-induced ECM deformations.** Interpenetrating networks (IPNs) were seeded with 0.5 µm fluorescent beads to visualize ECM mechanical deformations induced by cell-generated forces. CellTracker-labeled cells and the beads were imaged every 30 s using a Nikon 40× NA 1.15 oil-immersion objective. The 3D displacements of beads and cells were tracked individually in Imaris. Custom MATLAB codes were used to project displacement vectors onto XY, YZ, and ZX planes by interpolating bead displacements from discrete positions onto a continuous grid. Interpolation was performed using the natural neighbor method in MATLAB, which is based on geometric Voronoi tessellation. This parameter-free approach is strictly local, incorporates only the “natural neighbors” (beads with Voronoi cells directly affected), and passes exactly through the original data points, ensuring smooth interpolation while preserving physical consistency. Another MATLAB code was then used to compute 3D principal strains by (i) interpolating scattered bead displacements onto a regular 3D grid using the natural neighbor method, (ii) calculating the deformation tensor (displacement field gradient), and (iii) visualizing the displacement field.

**Statistics and reproducibility.** All experiments included 2-13 replicates from at least two independent preparations, with the exact sample size (n) and specific statistical test performed detailed in the corresponding figure captions. Unless stated otherwise, data are presented as mean ± standard error of the mean (SEM). The choice between parametric and non-parametric tests was determined after assessing data for normal distribution. Comparisons between two groups were performed using a two-tailed Student's t-test (for normal distributions) or a Mann-Whitney U test (for non-parametric data). Comparisons between multiple groups were performed using ordinary one-way ANOVA with Tukey's post-hoc analysis (for normal distributions) or a Kruskal-Wallis test with Dunn's post-hoc test (for non-parametric data). All statistical analyses were conducted using GraphPad Prism (Versions 9–11; GraphPad Software, La Jolla, CA, USA). P-values were considered statistically significant as follows: *$P < 0.033$, **$P < 0.002$, and ***$P < 0.001$.

### Supplementary Figures

### A. Pore size measurement of fresh tissues and an IPN

This section details the image analysis methodology used to quantify the pore sizes of fresh human tissues and an interpenetrating network (IPN) hydrogel. Figure S1 presents the raw and binarized images of fresh tissue samples along with their corresponding pore size measurements. The step-by-step procedure for determining the pore size distribution of an IPN hydrogel is illustrated in Figure S2. Note that essentially the same approach was applied to both the tissue and IPN pore size measurements.

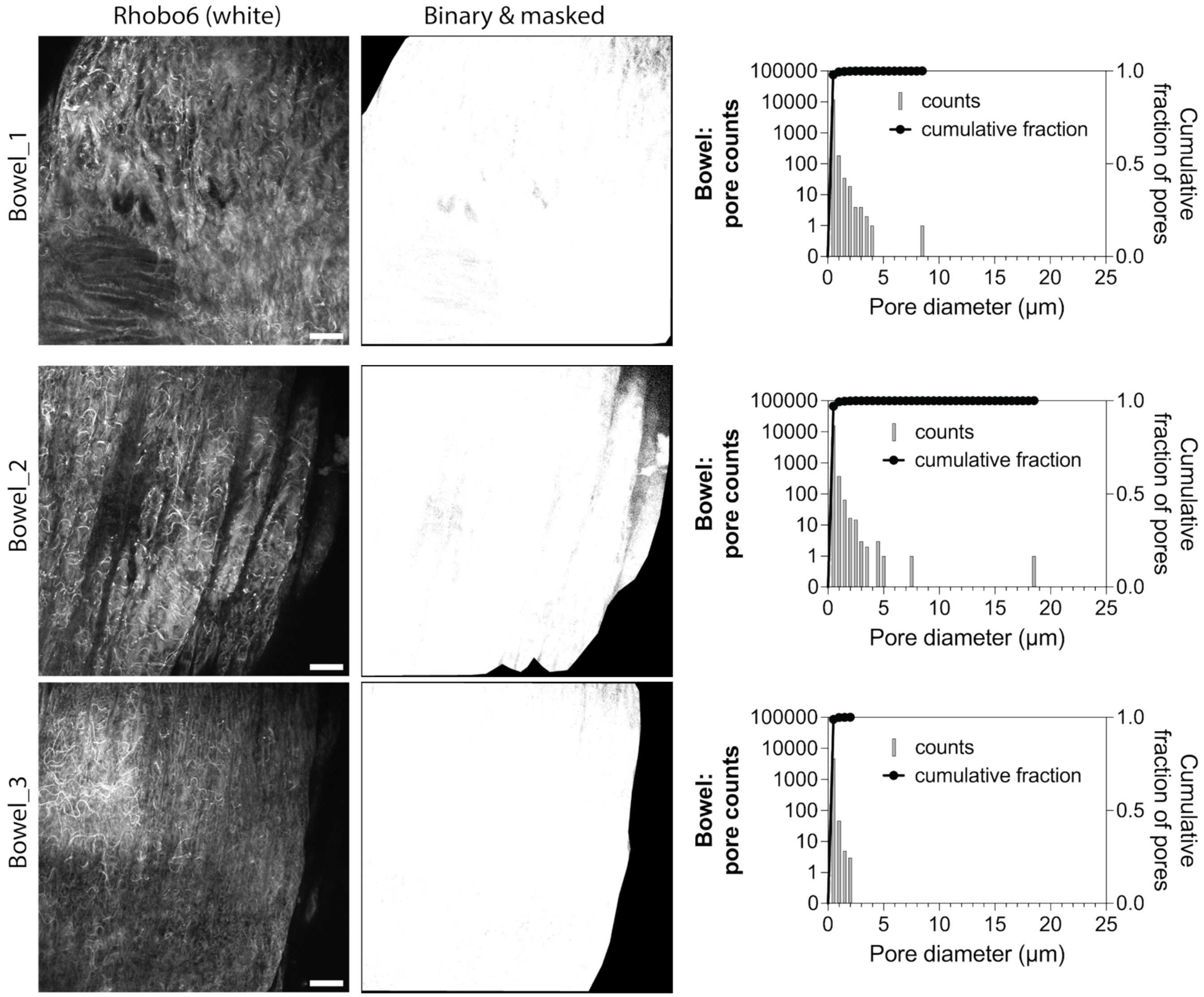

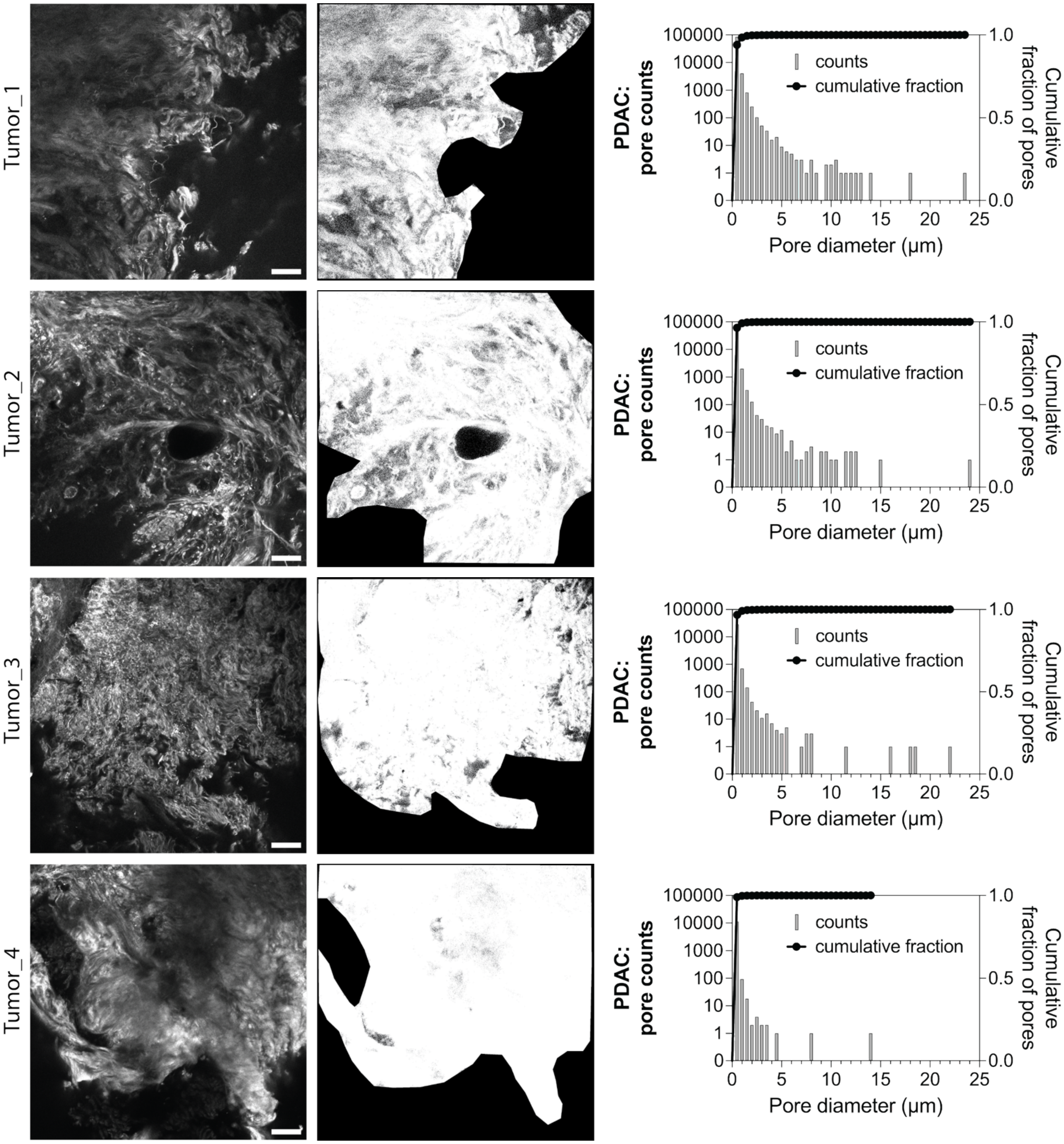

Tumor_1
Tumor_2
Tumor_3
Tumor_4
PDAC: pore counts
Cumulative fraction of pores
counts
cumulative fraction
Pore diameter (μm)
100000
10000
1000
100
10
1
0
0
5
10
15
20
25
1.0
0.5
0.0

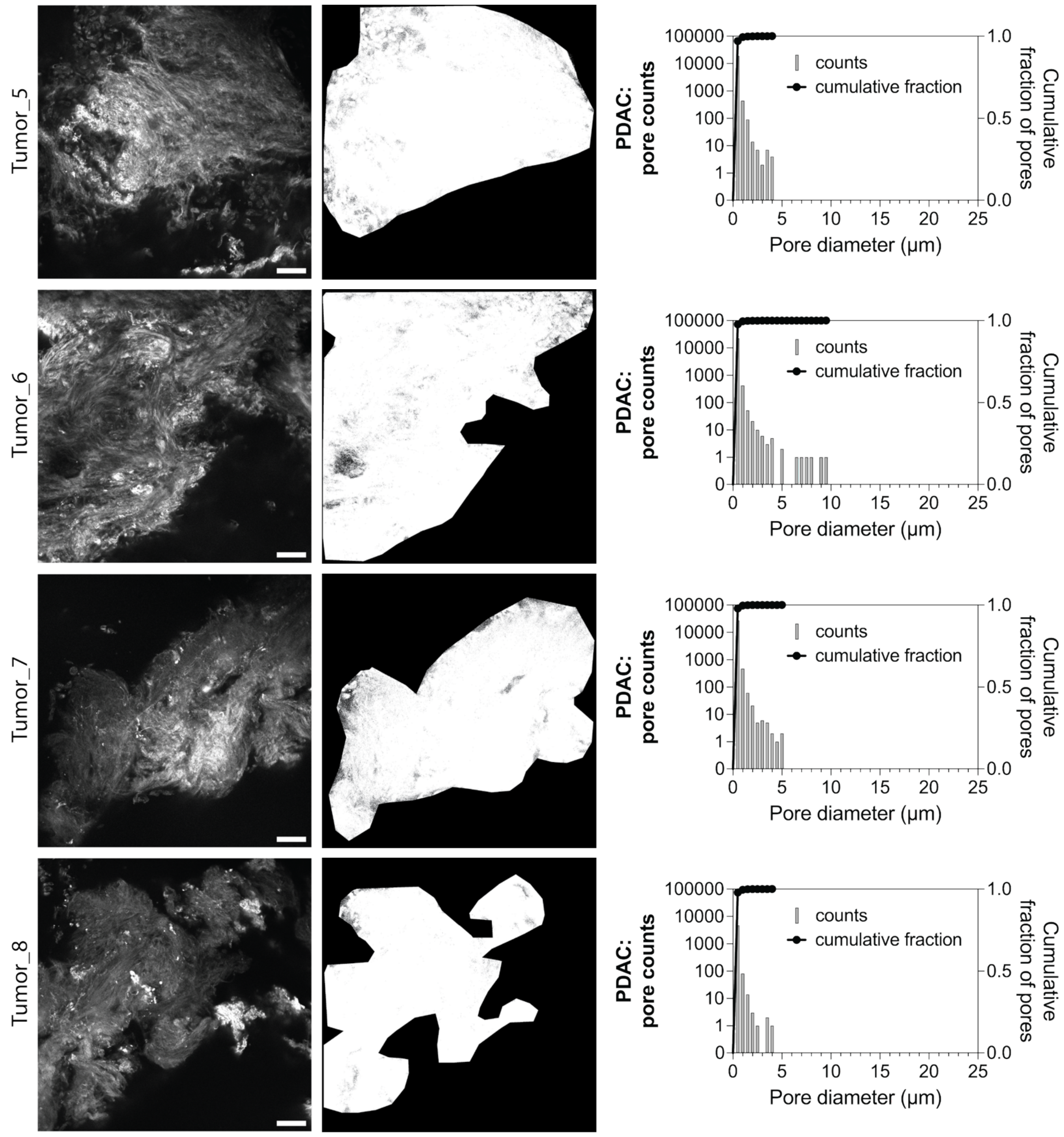
Tumor_5
Tumor_6
Tumor_7
Tumor_8
PDAC:
pore counts
Cumulative
fraction of pores
counts
cumulative fraction
100000
10000
1000
100
10
1
0
1.0
0.5
0.0
0
5
10
15
20
25
Pore diameter (μm)

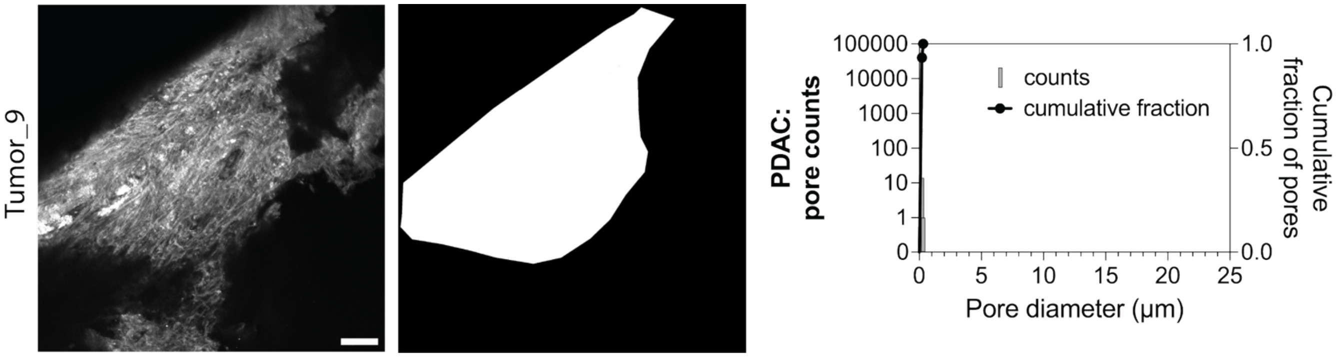


**Fig. S1. Pore size analysis of normal bowel and pancreatic ductal adenocarcinoma (PDAC) tissues.** The first column displays fluorescence micrographs of the extracellular matrix (ECM) in freshly excised human normal bowel (n=3) and PDAC (n=9) tissues, stained with Rhobo6. The second column presents corresponding binarized images, generated by masking empty spaces and thresholding based on background intensity. The third column shows pore counts and diameters measured from the dark regions identified from the corresponding second column images. Scale bar: 50 μm.

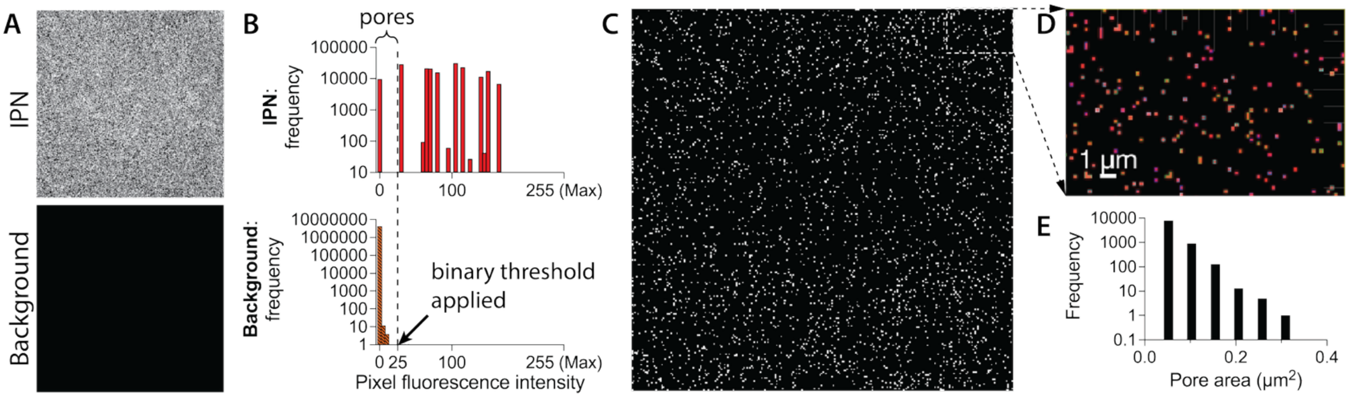


**Fig. S2. Pore size analysis in IPNs** (A) Fluorescence micrographs of (top) an IPN containing fluorescent alginate and (bottom) the media. (B) Fluorescence intensity distribution of all the pixels corresponding to the images of IPN and the media. Binary threshold was set above the maximum intensity of background. In the IPN image, the pixels exhibiting intensity lower than the threshold were regarded as pore area. (C) The binarized image of an IPN image. (D) The zoom-in image shows pores segmented by Imaris. (E) The number of pores is plotted as a function of its area.

#### B. Shear rheology

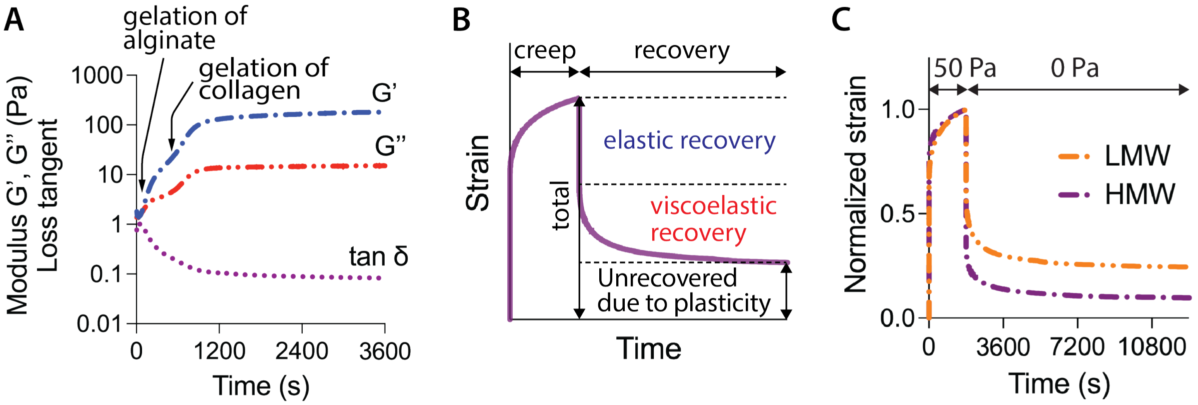


**Fig. S3. Shear rheology measurements on IPNs** (A) Representative oscillation test of an IPN. Two steep slopes of increasing moduli differentiated by a deflection curve in the early times respectively show the gelation of alginate followed by the gelation of type-1 collagen. The storage and loss moduli and loss tangent were recorded at the end of 1 hour-long test. (B) Schematic depicting the elastic, viscoelastic, and plastic (permanent) portions of a material response in a creep and recovery test. (C) Representative creep and recovery tests of IPNs.

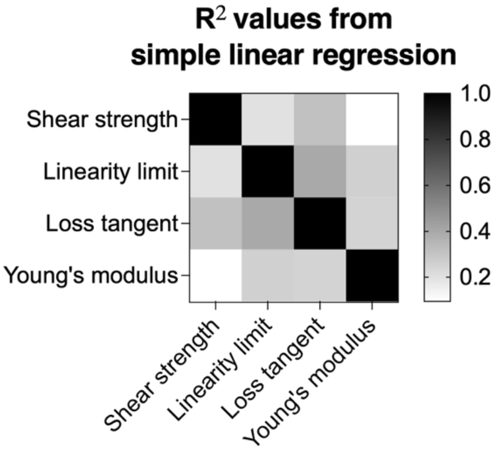


**Fig. S4. Heat map of coefficients of determination or $R^2$ values from simple linear regression.**

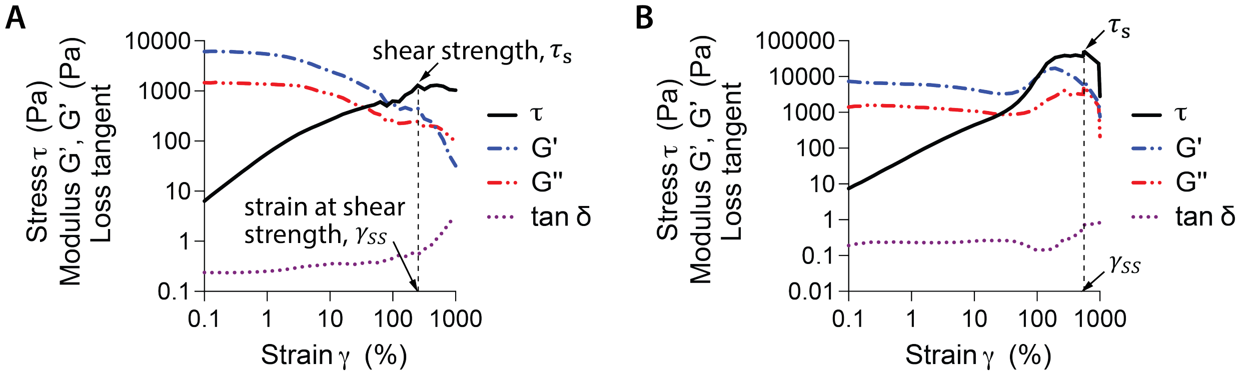


**Fig. S5. Representative stress-strain curve data for soft tissues**. These were measured from strain-controlled amplitude sweeps of (A) human normal pancreas (resected from a PDAC patient; biological replicates N=6 for the measurements of Young's modulus, loss tangent, shear strength shown in Fig. 1I) and (B) PDAC tissue (N=10). The shear strength ($\tau_S$) was defined as the maximum oscillatory stress at or before the point where the loss tangent reached 1. Once the loss tangent exceeded 1, indicating a transition from viscoelastic solid to viscoelastic liquid, subsequent stress values were not considered when determining shear strength, even if they surpassed earlier values. Such behavior was rarely observed, and only in soft normal tissues and particularly thin (<2 mm) samples, where we suspect that higher friction at large imposed strains required elevated stresses. Because the amplitude sweeps were strain-controlled, applying strains approaching 1000% could cause the sample to coil (like a twisted rope) or generate substantial friction, artificially increasing the measured stress. Note also that the storage and loss moduli measured during amplitude sweeps tend to be significantly higher due to the use of superglue, which stiffens tissue regions that bonds to the rheometer plates; these modulus values should therefore be interpreted with caution and were not used in our study.

### C. Full motility data of Jurkat and primary human T cells

In the main Figure 2, only subsets of the motility and correlation data are shown due to space constraints. Here, we provide the complete datasets for both Jurkat and primary human T cells.

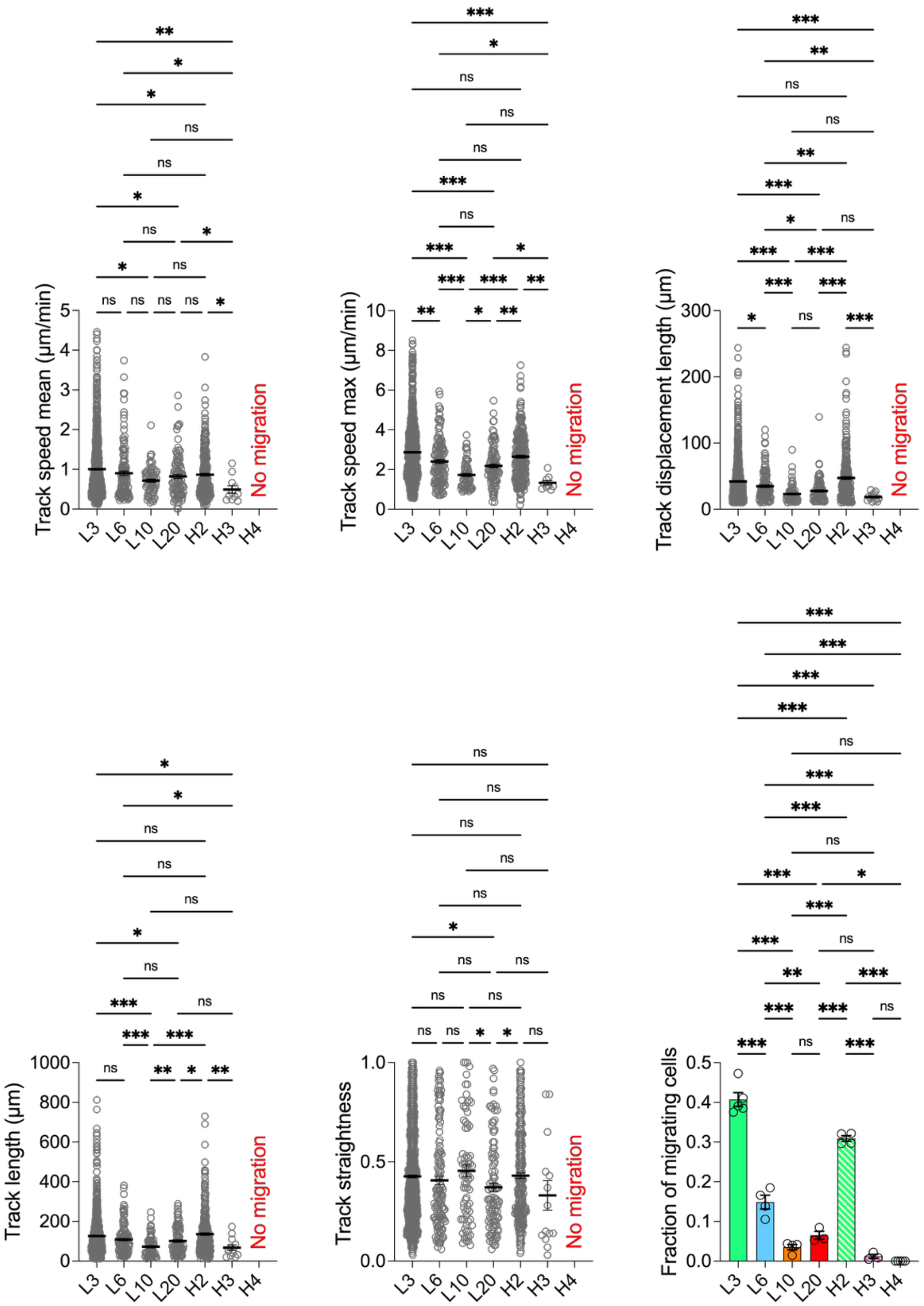


**Fig. S6. Motility metrics of Jurkat T cells in each IPN.** Data are shown for track speed mean, track speed max, track displacement length, track length, track straightness, and fraction of migrating cells among all tracked cells. Cells were classified as arrested if their track displacement length was <10 μm.

Motility metrics were quantified only for motile cells (track duration ≥10 min and track displacement length ≥10 μm). For each IPN condition, the total number of motile cell tracks (n), biological replicates (N), and experiments (E) are: L3 (n=1115, N=5, E=2), L6 (n=141, N=4, E=2), L10 (n=78, N=4, E=2), L20 (n=108, N=3, E=2), H2 (n=285, N=4, E=2), H3 (n=14, N=3, E=2). For H4 IPN, where all cells were arrested, 281 cells were tracked across 5 replicates from 3 experiments. *$P < 0.033$, **$P < 0.002$, and ***$P < 0.001$ by one-way ANOVA followed by Tukey's post hoc test (fraction of migrating cells) or Kruskal-Wallis followed by uncorrected Dunn's test (the other motility metrics). All data are shown as means ± SEM.

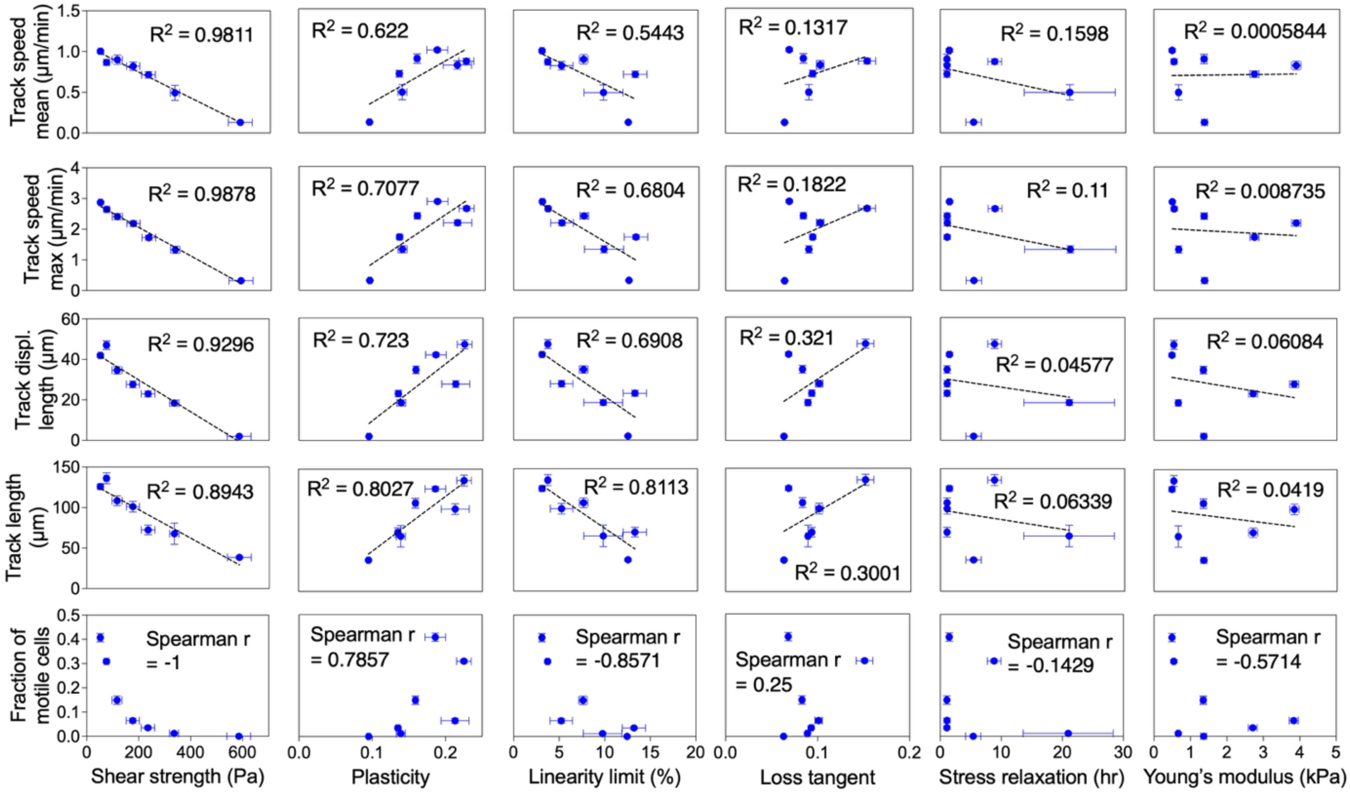


**Fig. S7. Correlation of the Jurkat T cell motility and the IPN mechanical properties**. Data was assessed using Spearman's rank correlation and simple linear regression.

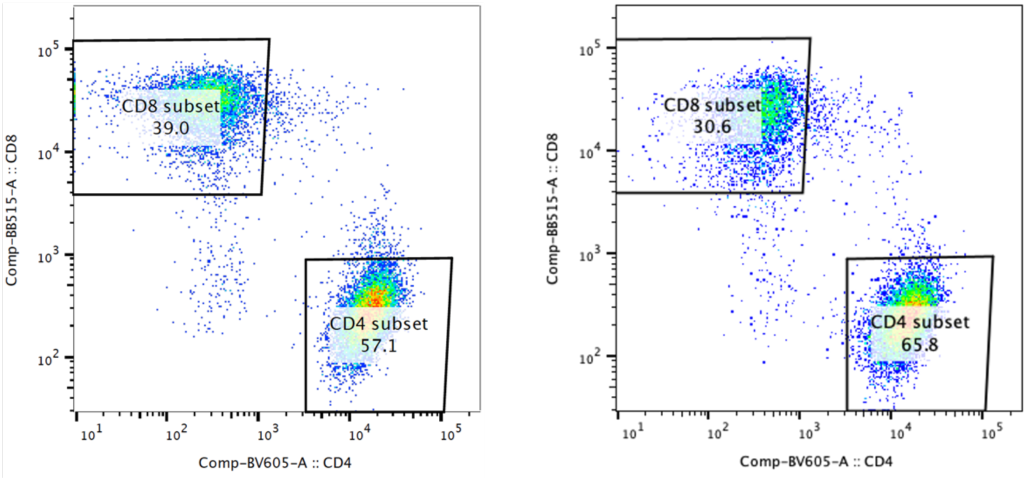


**Fig. S8. Flow cytometry analysis for two experiments.** These data confirm that primary human T cells used for our research consist of ~60% CD4+ and ~40% CD8+ subtypes.

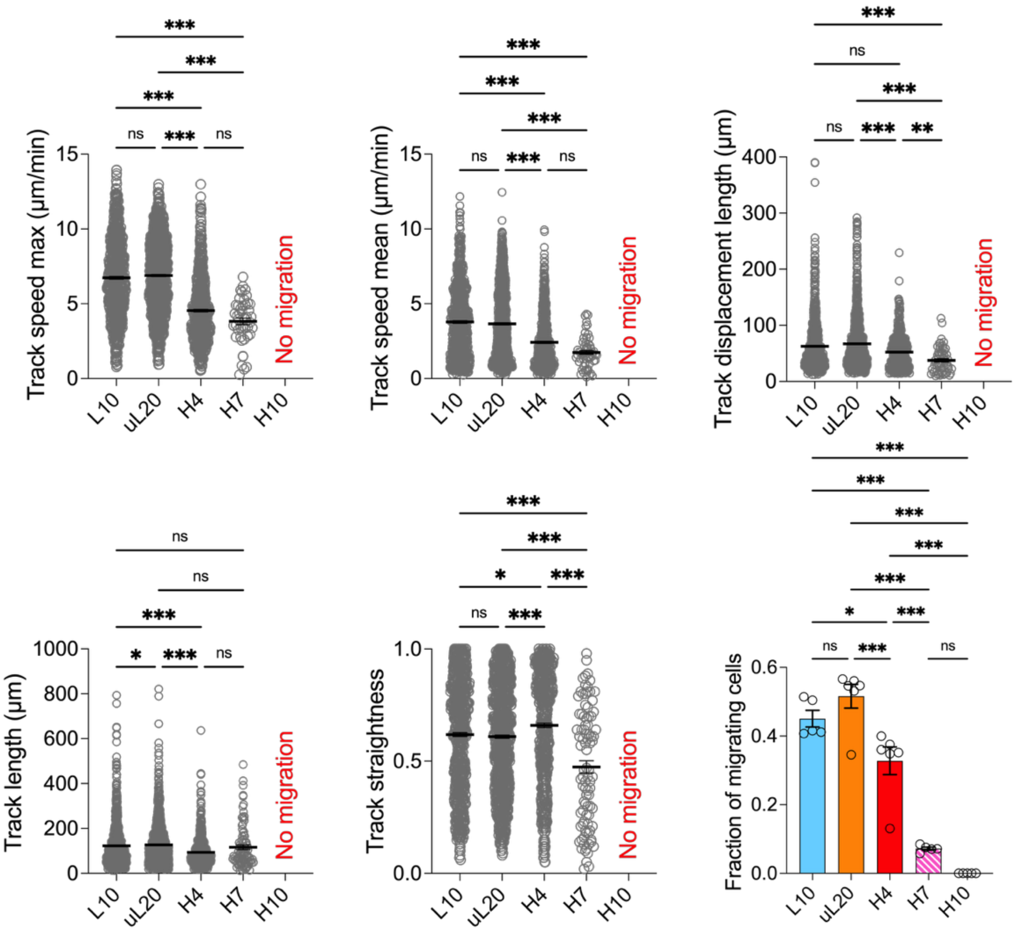


**Fig. S9. Motility metrics of primary T cells measured per each IPN.** Data are shown for track speed mean, track speed max, track displacement length, track length, track straightness, and fraction of migrating cells of all tracked cells. Cells were classified as arrested if their track displacement length fell below a set threshold. This threshold, ranging from 8 to 20 µm, was determined for each dataset individually to account for any residual system drift after computational correction. Motility metrics were quantified only for motile cells (track duration ≥5 min and track displacement length exceeding the 8–20 µm threshold). For each IPN condition, the total number of motile cell tracks (n), biological replicates (N), and experiments (E) are: L10 (n=782, N=5, E=2), uL20 (n=1115, N=6, E=2), H4 (n=625, N=6, E=3), H7 (n=91, N=5, E=2). For H10 IPN, where all cells were arrested, 1217 cells were tracked across 5 replicates from 2 experiments. *$P < 0.033$, **$P < 0.002$, and ***$P < 0.001$ by One-way ANOVA followed by Tukey's (fraction of migrating cells) and Kruskal-Wallis & uncorrected Dunn's test (the other motility metrics) post hoc analysis. All data are shown as means ± SEM.

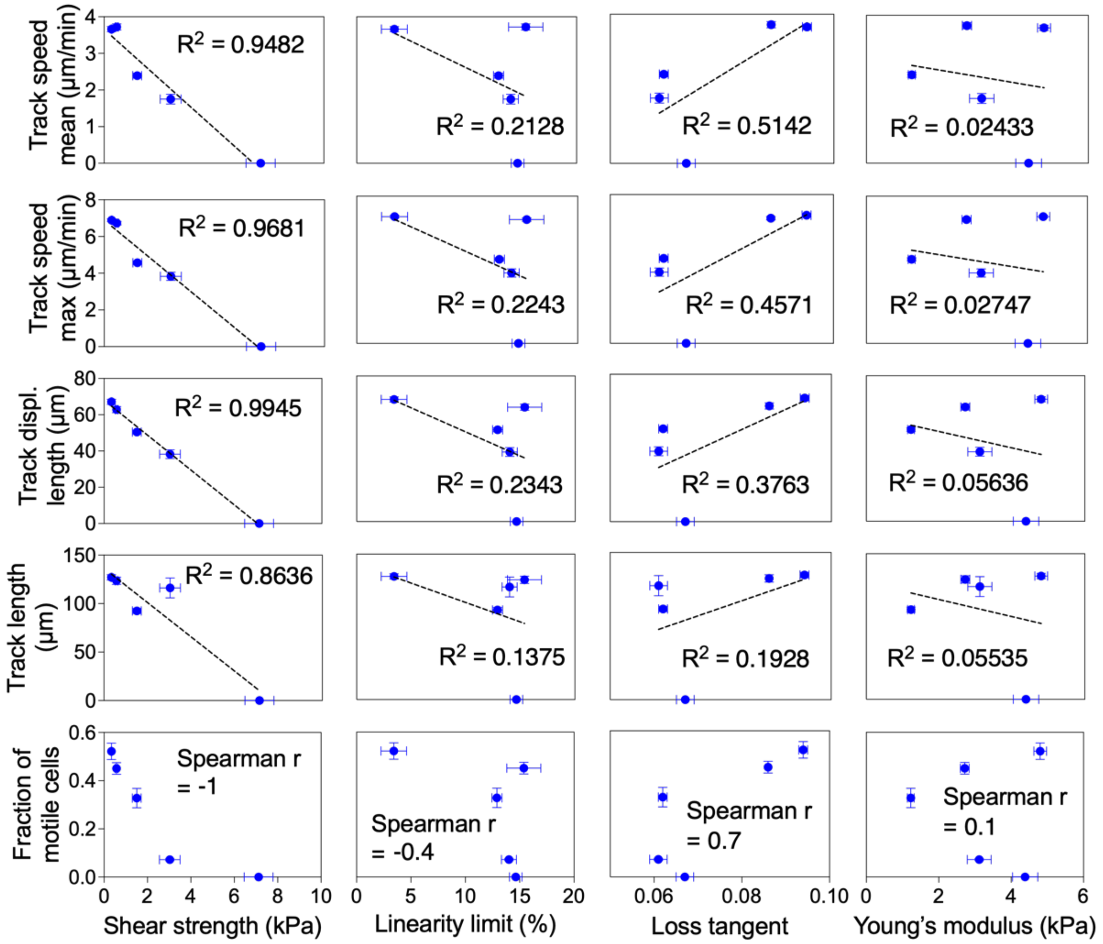


**Fig. S10. Correlation of the primary T cell motility and the IPN mechanical properties.** Data were assessed using simple linear regression and Spearman's rank correlation.

#### D. Cell motility and viability in pharmacological inhibition studies

In the Figure 3, only subsets of the motility data are shown due to space constraints. Here, we provide additional datasets.

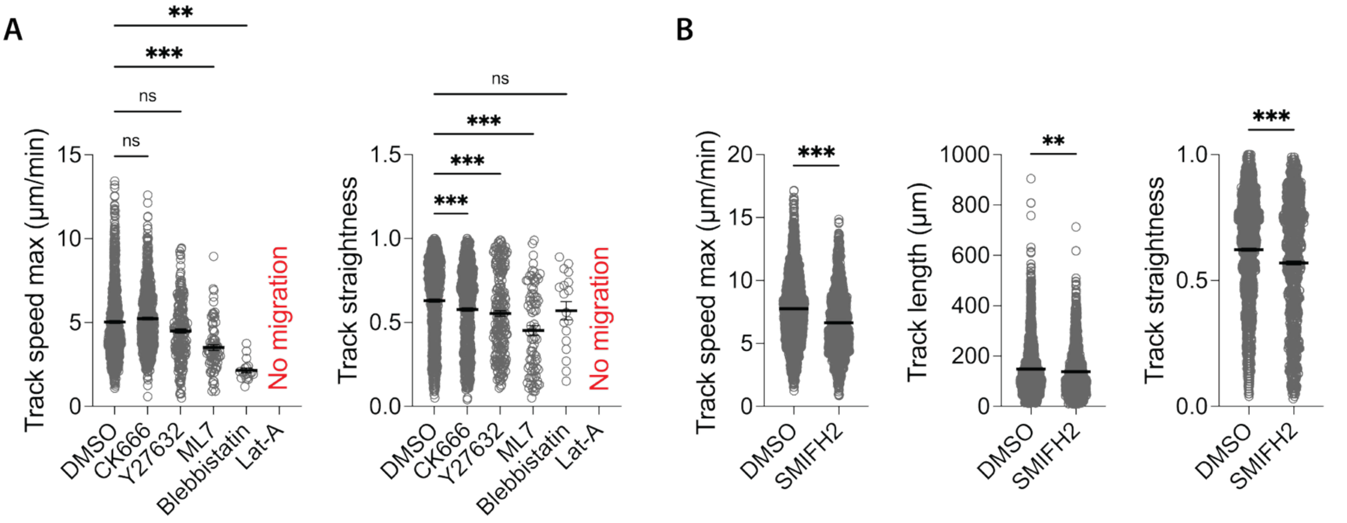


**Fig. S11. Additional data showing the effect of pharmacological inhibition on cell motility.** (A) Kruskal-Wallis and corrected Dunn's test; n≥615 for each condition; N=4 replicates; 2 experiments. (B) Unpaired, two-tailed Mann-Whitney U test; n≥1619 for each condition; N=4 replicates; 2 experiments. *P < 0.033, **P < 0.002, and ***P < 0.001. All data are shown as means ± SEM.

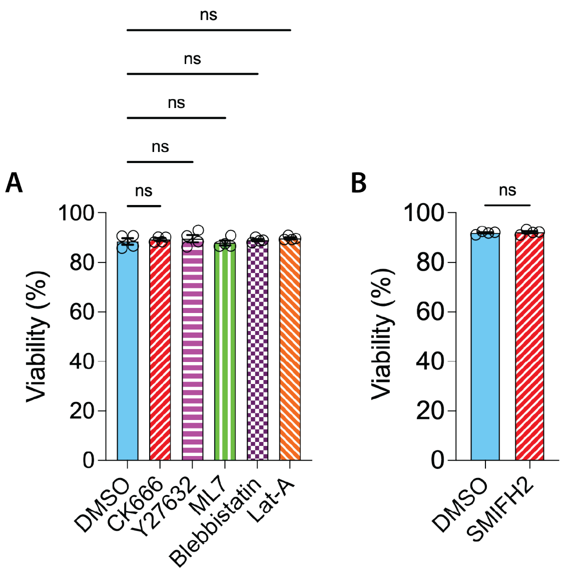


**Fig. S12. Effect of pharmacological inhibition on cell viability.** (A) One-way ANOVA followed by Tukey's post hoc analysis; Total number of cell counts: n=689 (DMSO), 439 (CK666), 471 (Y27632), 480 (ML7), 701 (Bleb), 569 (Lat-A); N=4 replicates; 2 experiments. (B) Unpaired t test; Total number of cell counts: n=1207 (DMSO), 981 (SMIFH2); N=4 replicates; 2 experiments. *P < 0.033, **P < 0.002, and ***P < 0.001. All data are shown as means ± SEM.

### E. Bead displacement assays

In the Figure 4, only a subset of the bead displacement assay data is shown due to space constraints. Substantial differences in size, shape, and migration speed from cell to cell preclude generation of an accurate averaged traction strain map. Thus, we provide additional datasets demonstrating the traction strain patterns for other migrating cells.

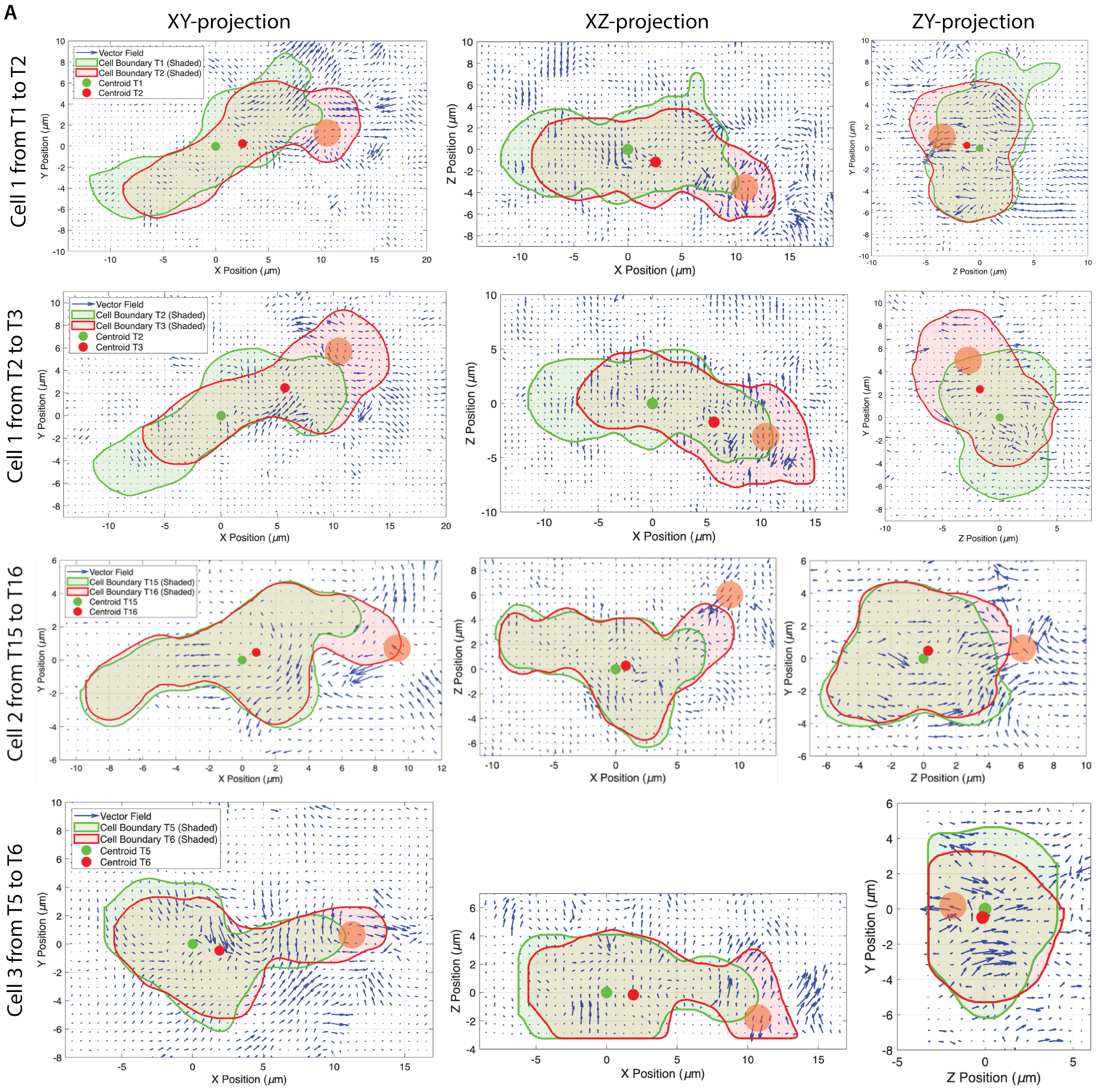

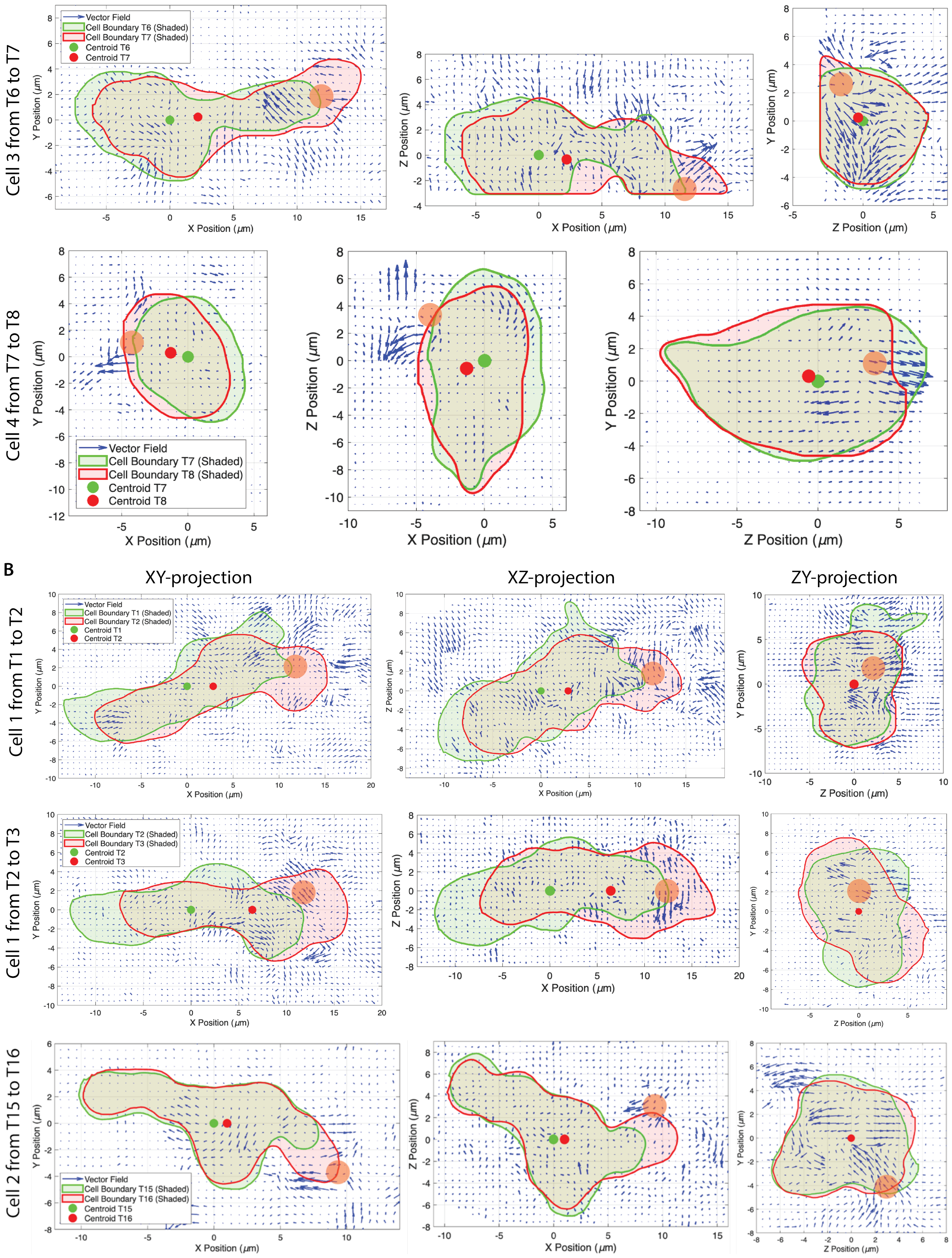
Cell 3 from T6 to T7
Vector Field
Cell Boundary T6 (Shaded)
Cell Boundary T7 (Shaded)
Centroid T6
Centroid T7
X Position (μm)
Y Position (μm)
Z Position (μm)
Cell 4 from T7 to T8
Vector Field
Cell Boundary T7 (Shaded)
Cell Boundary T8 (Shaded)
Centroid T7
Centroid T8
B
XY-projection
XZ-projection
ZY-projection
Cell 1 from T1 to T2
Vector Field
Cell Boundary T1 (Shaded)
Cell Boundary T2 (Shaded)
Centroid T1
Centroid T2
Cell 1 from T2 to T3
Vector Field
Cell Boundary T2 (Shaded)
Cell Boundary T3 (Shaded)
Centroid T2
Centroid T3
Cell 2 from T15 to T16
Vector Field
Cell Boundary T15 (Shaded)
Cell Boundary T16 (Shaded)
Centroid T15
Centroid T16

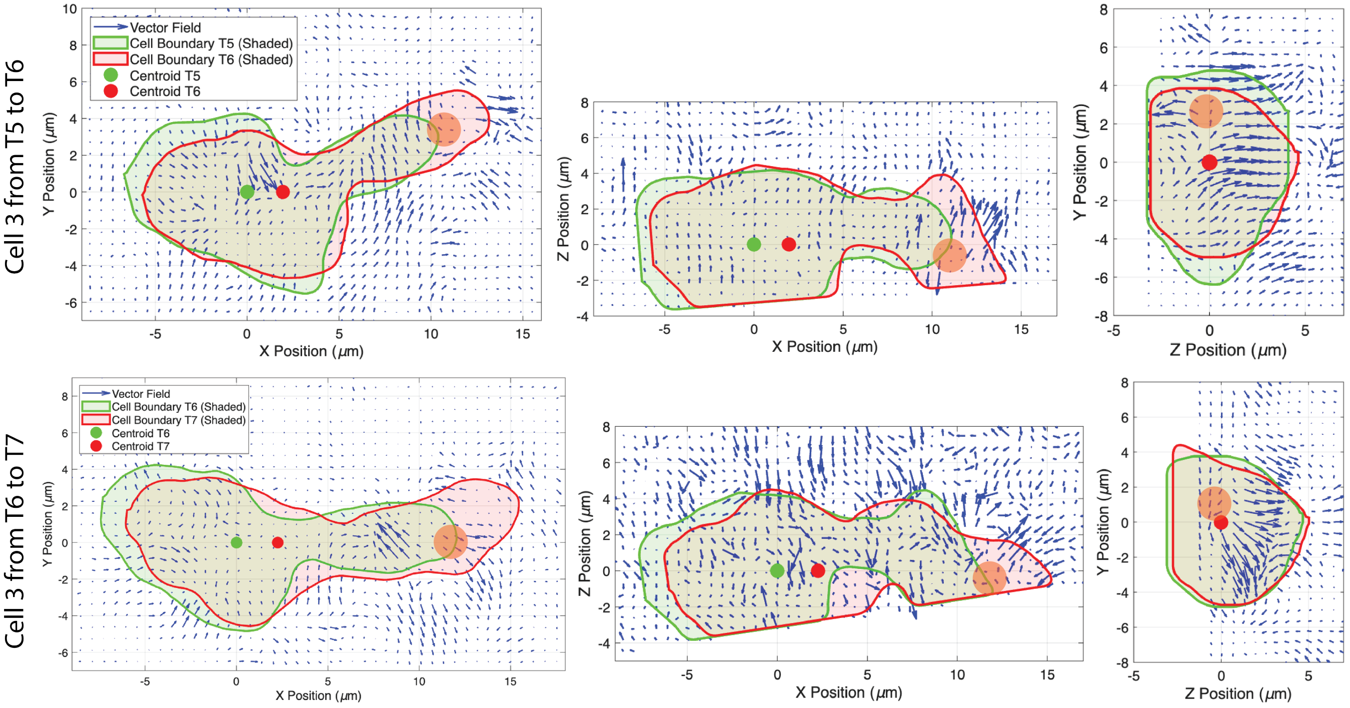


**Fig. S13. 2D projection maps of bead displacements around a migrating cell.** (A) Bead displacements in a fixed coordinate system. 3D bead displacements, tracked in 30-second intervals using Imaris, were projected onto 2D (XY, XY, ZY) planes. Cell boundaries, determined from Imaris snapshots with a custom MATLAB code, and centroids at the preceding ($T_i$) and subsequent ($T_{i+1}$) time points are overlaid. The orange circle indicates the estimated location of the cell's leading edge, identified by the convergence point of the displacement vectors and its proximity to the region of maximum principal strain. This fixed-frame view is useful for visualizing changes in the cell's overall migration trajectory. (B) Bead displacements in a rotated coordinate system. To create a "head-on" perspective in the ZY plane that captures radially divergent strain originating from the leading pole, the displacement field was computationally rotated. The rotation aligns the cell's migration vector—defined from the centroid at $T_i$ to $T_{i+1}$—with the positive x-axis (1,0,0). It should be noted that this is an approximation, as the centroid displacement vector may not perfectly align with the instantaneous migration direction, especially during abrupt turns. Nevertheless, this rotated view effectively captured some radially divergent matrix deformation—which were often not clearly visible in the fixed coordinates—demonstrating that the "breaststroke" motion originates from the leading edge and undergoes radially divergent, rearward movement. A total of four cells were tracked, and the images shown are from representative time points.

### Supplementary Tables

| | 7 IPN formulations for Jurkat cells | | | | | | | 5 IPNs for primary T cells | | | | |
|---|---|---|---|---|---|---|---|---|---|---|---|---|
| **Notation** | **L3** | **L6** | **L10** | **L20** | **H2** | **H3** | **H4** | **L10** | **uL20** | **H4** | **H7** | **H10** |
| $Ca^{2+}$ (mM) | 3 | 6 | 10 | 20 | 2 | 3 | 4 | 10 | 20 | 4 | 7 | 10 |
| alg MW | L | L | L | L | H | H | H | L | uL | H | H | H |
| alg (mg/mL) | 6.0 | 6.0 | 6.0 | 6.0 | 6.0 | 6.0 | 6.0 | 6.0 | 6.0 | 6.0 | 6.0 | 6.0 |
| col-1 (mg/mL) | 1.5 | 1.5 | 1.5 | 1.5 | 1.5 | 1.5 | 1.5 | 1.5 | 1.5 | 1.5 | 1.5 | 1.5 |

**Table. S1. Composition and notation of IPN formulations.** Sets of seven and five IPN formulations were developed respectively to study Jurkat and primary T cell migration. The notation system defines each formulation by alginate type and crosslinker concentration: 'L' denotes low-molecular-weight (LMW) alginate, 'uL' denotes unfiltered, unlyophilized LMW alginate, and 'H' denotes high-molecular-weight (HMW) alginate. The number indicates the calcium crosslinker concentration in mM (e.g., L3 is an IPN of LMW alginate with 3 mM calcium). All IPNs contained the same concentrations of alginate (6.0 mg/mL) and collagen-I (1.5 mg/mL) to ensure consistent nanoporosity and adhesive ligand density.

**Supplementary Movie captions**

**Movie S1. Jurkat T cell migration through a nanoporous IPN.**

**Movie S2. Jurkat T cells show high motility in a low-strength (L6) IPN**

**Movie S3. Jurkat T cell migration is arrested in a high-strength (H4) IPN**

**Movie S4. Primary human T cell migration through a nanoporous IPN.** Frames per second was set such that the play speed is 100 times faster than the real.

**Movie S5. Primary human T cells show high motility in a low-strength (uL20) IPN**

**Movie S6. Primary human T cells show limited motility in an intermediate-strength (H7) IPN**

**Movie S7. Primary human T cell migration is arrested in a high-strength (H10) IPN.** Time-lapse imaging over 9 hours shows that the cells remain completely immobilized within the matrix.

**Movie S8. Brightfield live-imaging reveals that primary human T cell shows breaststroke-like membrane movement during migration.**

**Movie S9. Fluorescence live-imaging reveals the actin dynamics of the primary human T cell migrating through an IPN.**

**Movie S10. Individual beads around a migrating cell were individually tracked via time-lapse confocal microscopy and Imaris analysis.**

**Movie S11. Time-lapse confocal microscopy of primary T cell migration in a fluorescent-alginate IPN, shown as a three-channel overlay of the cells, alginate, and collagen-1 (via reflectance).**

**Movie S12. Time-lapse confocal microscopy of primary T cell migration in a fluorescent-alginate IPN, shown as a two-channel overlay of the cells and alginate.**
